# The Splice Index as a prognostic biomarker of strength and function in myotonic dystrophy type 1

**DOI:** 10.1101/2024.07.10.602610

**Authors:** Marina Provenzano, Kobe Ikegami, Kameron Bates, Alison Gaynor, Julia M. Hartman, Aileen S. Jones, Amanda Butler, Kiera N. Berggren, Jeanne Dekdebrun, Man Hung, Dana M. Lapato, Michael Kiefer, Charles Thornton, Nicholas E. Johnson, Melissa A. Hale, the Myotonic Dystrophy Clinical Research Network (DMCRN)

## Abstract

Myotonic dystrophy type 1 (DM1) is a slowly progressive, multisystemic disorder caused by a CTG repeat expansion in the *DMPK* 3’UTR that leads to global dysregulation of alternative splicing. Here, we employed a composite RNA splicing biomarker called the Myotonic Dystrophy Splice Index (SI), which incorporates 22 disease-specific splice events that sensitively and robustly assesses transcriptomic dysregulation across the disease spectrum. Targeted RNA sequencing was used to derive the SI in 95 muscle biopsies of the tibialis anterior collected from DM1 individuals with baseline (n = 52) and 3-months (n = 37) outcomes. The SI had significant associations with timepoint matched measures of muscle strength and ambulation, including ankle dorsiflexion strength (ADF) and 10-meter run/fast walk speed (Pearson *r* = -0.719 and -0.680, respectively). Linear regression modeling showed that the combination of baseline ADF and SI was predictive of strength at 3-months (adjusted R^2^ = 0.830) in our cohort. These results indicate the SI can reliably capture the association of disease-specific RNA mis-splicing to physical strength and mobility and may be predictive of future function.

## Introduction

Myotonic Dystrophy type 1 (DM1) is the most common form of adult-onset muscular dystrophy with an estimated gene frequency of 1:2100^1^. This autosomal dominantly inherited disorder is characterized by a core triad of myotonia (i.e. difficulty relaxing muscles post contraction), progressive distal muscle weakness, and early onset cataracts. Beyond these characteristic symptoms, DM1 is a multisystemic disorder affecting nearly every organ in the body including the central nervous, gastrointestinal, and cardiovascular systems^2^. The impact on these organ systems results in the presentation of daytime sleepiness, cardiac arrhythmias, and respiratory failure. This complex presentation leads to a phenotypically diverse population with variable symptomatic presentation and progression.

DM1 is caused by an expanded CTG repeat tract (CUG_exp_) in the 3’ untranslated region of the *DMPK* gene^3–5^. (CUG)_exp_ RNAs transcribed from this locus are toxic due to their propensity to sequester RNA binding proteins, most notably muscleblind-like (MBNL)^6–8^. This alternative splicing factor is a key driver of fetal to adult RNA isoform transitions during development and its sequestration leads to widespread dysregulation of alternative splicing and retention of mRNA transcripts encoding non-functional fetal isoforms of the associated gene products^9–13^. Mis-splicing of select MBNL-dependent events have been linked to disease symptoms including insulin resistance (*INSR*), myotonia (*CLCN1*), and cardiac arrhythmia (*SCN5A*) in DM1 cell and mouse models^14–20^. Prior work has demonstrated that these are titratable events whereby a dose-response relationship is observed between changes in RNA splicing and the relative sequestration of MBNL1^21,22^. This dose-dependent relationship is event-specific whereby individual RNA splice events are more or less sensitive to functional concentrations of MBNL associated with the spectrum of DM1 disease severity^21,22^. In concordance with these observations, RNA splicing events are differentially rescued in DM1 pre-clinical models across a therapeutic dose-range and have been shown to differentially respond to interventional knockdown of the *DMPK* transcript in a human clinical trial^23–28^. These findings have been mirrored in evaluation of mis-splicing within a group of DM1 individuals where select events had stronger or weaker correlations with manual muscle testing of the ankle dorsiflexion^29^.

Given its relevance to disease pathology and correlation with physical function in DM1, RNA splicing of MBNL-dependent, disease-associated events is a promising biomarker for pre-clinical and clinical evaluation of therapeutic engagement. However, in isolation, no single event has been shown to capture the range of molecular and phenotypic variability observed in DM1 individuals making it difficult to match the degree of mis-splicing of an isolated RNA event to the status of overall physical function in an individual^29^. Previous work has sought to develop a composite splicing measure that could capture the range of phenotypic severity observed. In DM1 mouse models a composite measure of 35 splice events showed a graded response to decrements of Mbnl or increasing (CUG)_exp_ load^27^. Early efforts to identify human candidate biomarkers utilized a measure known as [MBNL]_inferred_ which utilizes a small collection of RNA splicing events highly predictive of global mis-splicing changes in DM1 tibialis anterior muscle, to estimate intracellular levels of non-sequestered MBNL^21,30^. However, these studies have been limited by a lack of skeletal muscle biospecimens from a large, well-characterized cohort inclusive of longitudinal sampling to comprehensively profile the complete spectrum of RNA mis-splicing and to assess the dynamics of mis-splicing over time.

Here, we generated a total RNA sequencing (RNAseq) dataset encompassing 95 skeletal muscle transcriptomes from an extensively phenotyped cohort of 58 DM1 participants, a subset of whom provided longitudinal biopsies at baseline and a 3-month follow up visit. This dataset was utilized to assess the validity of a composite RNA splicing biomarker called the Myotonic Dystrophy Splice Index (SI). Leveraging high-resolution, targeted amplicon RNAseq of this 22-event panel, we demonstrate that the SI accurately assesses and normalizes the relative degree of skeletal muscle splicing dysregulation across the DM1 disease spectrum, is dynamic and sensitive to changes over time, and displays strong correlations with multiple measures of concurrent and future muscle performance. We also evaluate the ability of the SI to stratify mild, moderate, and severe cases of DM1 and leverage our longitudinally sampled sub-cohort to assess the utility of this biomarker to predict changes in physical function over time. This molecular marker provides a useful tool to capture the spectrum of DM1 disease severity and progression and may serve as a biomarker of target engagement in clinical trials.

## Results

### Phenotypically characterized DM1 participant cohort displays representative heterogenous distribution of global splicing dysregulation across the spectrum of disease severity

To facilitate DM1 biomarker development, we assembled a cohort of 58 adult-onset DM1 participants aged 21 to 70 years (**Fig 1A**, **Sup Table 1**). In this cohort, a total of 95 tibialis anterior (TA) biopsies were available for analysis with 35 individuals providing both baseline (BL) and 3-month (3M) longitudinal samples with corresponding timepoint-matched assessments of muscle performance. Cross-sectionally, this cohort displayed a wide range of muscle strength and performance as measured by Quantitative Muscle Testing (QMT) and Timed Motor Tests (TMT). There was broad variability in the degree of myotonia (the inability to relax muscles post contraction), as measured by the video hand opening time (vHOT) (**Fig 1B & Sup Fig 1A**).

**Figure 1:**
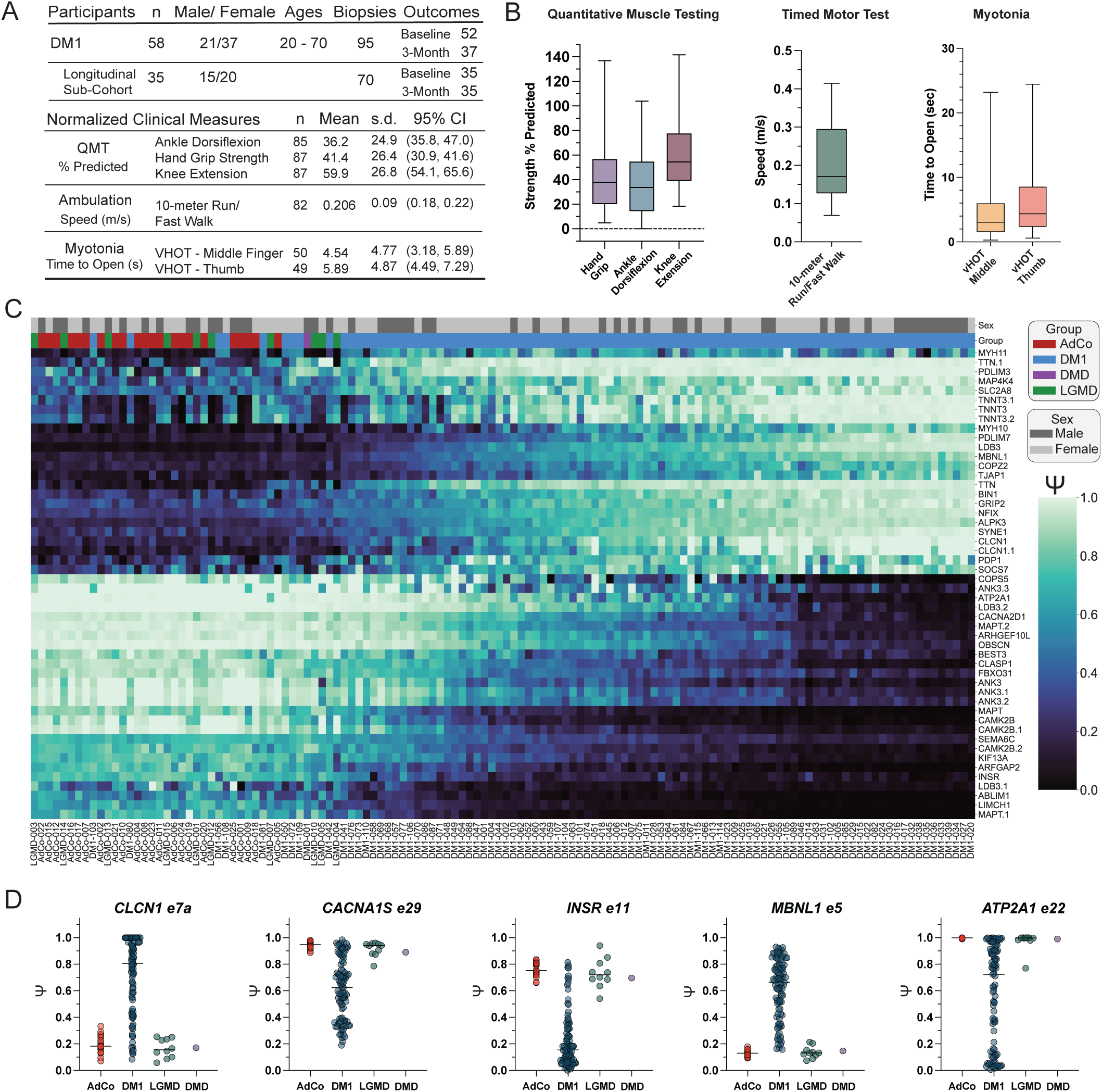
DM1 participant cohort displays broad spectrum of RNA splicing dysregulation assessed by total RNAseq. a) DM1 participant demographic information including sample size, age at biopsy, and sex distribution in complete cross-sectional and longitudinal cohort subset. The mean normalized performance on clinical outcome measures for the complete cross-sectional cohort are reported along with sample size, standard deviation (s.d.) and 95% confidence intervals (95% CI). Additional demographic information of all samples used in this report are provided in Sup Table 1. b) Box and whisker plot of outcome measure performance in cross-sectional cohort where line represents median performance and whiskers extend to max and minimum values. c) Heatmap displaying estimated percent spliced in (Ψ) of top 50 significantly dysregulated skipped exon (SE) events between DM1 versus unaffected adult controls (AdCo) and disease control reference groups (DMD & LGMD) subjected to total RNA-seq (|ΔΨ| ≥ 0.1, FDR ≤ 0.05). Both rows (SE events) and columns (individual samples) were subjected to hierarchical clustering. Sample group and sex are annotated above the heatmap and individual Subject IDs are reported below. d) Ψ values for specific SE events in all sample groups. Bar represents median values in each group.

To obtain a comprehensive view of RNA mis-splicing we performed total RNAseq on all available DM1 samples (n = 95) and evaluated global RNA splicing dysregulation. Additional muscle biopsies were included as reference groups from unaffected adults (AdCo, n = 22) and subjects with muscular dystrophies whereby the pathogenic mechanism is not predicted to cause RNA mis-splicing, including Duchenne muscular dystrophy (DMD) and limb-girdle (LGMD) types R1/2A, R3/2D, R7/2G, and R12/2L (n = 11) (**Sup Table 1**)^31–35^ While clinical outcomes were not available for these unaffected and disease-control participant sub-groups, they are included in this analysis to assess relative splicing dysregulation across the assembled DM1 cohort and disease specificity of observed mis-splicing. Inclusion levels (Ψ) for skipped exon (SE) splicing events were calculated via comparison of all DM1 subjects versus unaffected individuals and disease controls.

Principal component analysis of significantly mis-spliced events (|ΔΨ| ≥ 0.1, FDR ≤ 0.05, n = 946 SE events) illustrated the expected distribution; while unaffected individuals clustered tightly, all DM samples were distributed evenly across the first principal component consistent with the well-established spectrum of splicing dysregulation observed in other studies (**Sup Fig 1B**)^21,30^. The heterogeneity of mis-splicing within the full DM1 cohort is further demonstrated by visualization the top 50 most differentially spliced events (**Fig 1C** & **Sup Table 2**). While controls had minimal variability in Ψ, DM1 subjects possessed the full spectrum of splicing dysregulation; some individuals exhibited splicing patterns similar to AdCo and others exhibited maximal patterns of mis-splicing. All disease-controls clustered tightly with AdCo samples consistent with the predicted lack of significant RNA mis-splicing dysregulation associated with these muscular dystrophy sub-types (**Fig 1C** & **Fig 1B**) ^31–35^. The full spread of splicing dysregulation in this DM1 cohort is pronounced via visualization of select disease-specific events, including *CLCN1* e7a, *CACNA1S e29*, *INSR* e11, *ATP2A1 e22*, and *MBNL1* e5 (**Fig 1D**)^14–18,20,36–38^. In total, this DM1 cohort encapsulates a comprehensive spectrum of disease-specific, global splicing dysregulation and a range of skeletal muscle phenotypes. This combination positions this dataset as an effective biomarker assessment cohort which captures a representative snapshot of the range of molecular disease pathology.

### Selection and evaluation of splice events for inclusion in composite RNA splicing biomarker panel

While total RNAseq is a powerful discovery tool to comprehensively capture and describe global splicing dysregulation in DM1, ultra-deep sequencing coverage is required for precise quantification of exon inclusion. Computational estimation of Ψ inherently relies on sequence assembly and adequate splice-site junction coverage. As such, short paired-end sequencing like that typically utilized for transcriptomic analysis has reduced power for detecting of target splice events in transcripts that show low abundance or complex patterns of splice site utilization. If a subset of splicing events can distill patterns of global splicing dysregulation, targeted amplicon RNAseq utilizing extended read-lengths can be a more efficient tool to sensitively identify changes in RNA splicing. Previously, total RNAseq was used to identify 35 candidate splice events for inclusion in a targeted RNAseq panel for development of a species-specific composite splicing metric in DM1 mouse models^27^.

Building on these past efforts, we leveraged our well-stratified DM1 cohort to determine whether a subset of selected splice events can adequately capture the full range splicing dysregulation across a broad spectrum of disease severity. We utilized a panel of events that displayed widespread, significant changes in Ψ across our DM1 muscle transcriptomes compared to unaffected controls (**Fig 2A**)^28^. The panel incorporates 22 events as prior statistical modeling has indicated improved accuracy in estimating changes in MBNL activity using between 20 – 30 events^21^. Note that all events in the panel showed strong DM1 disease-specificity with limited mis-splicing observed in LGMD and DMD subjects and minimal variability in estimated Ψ within unaffected controls (**Sup Fig 2**).

**Figure 2:**
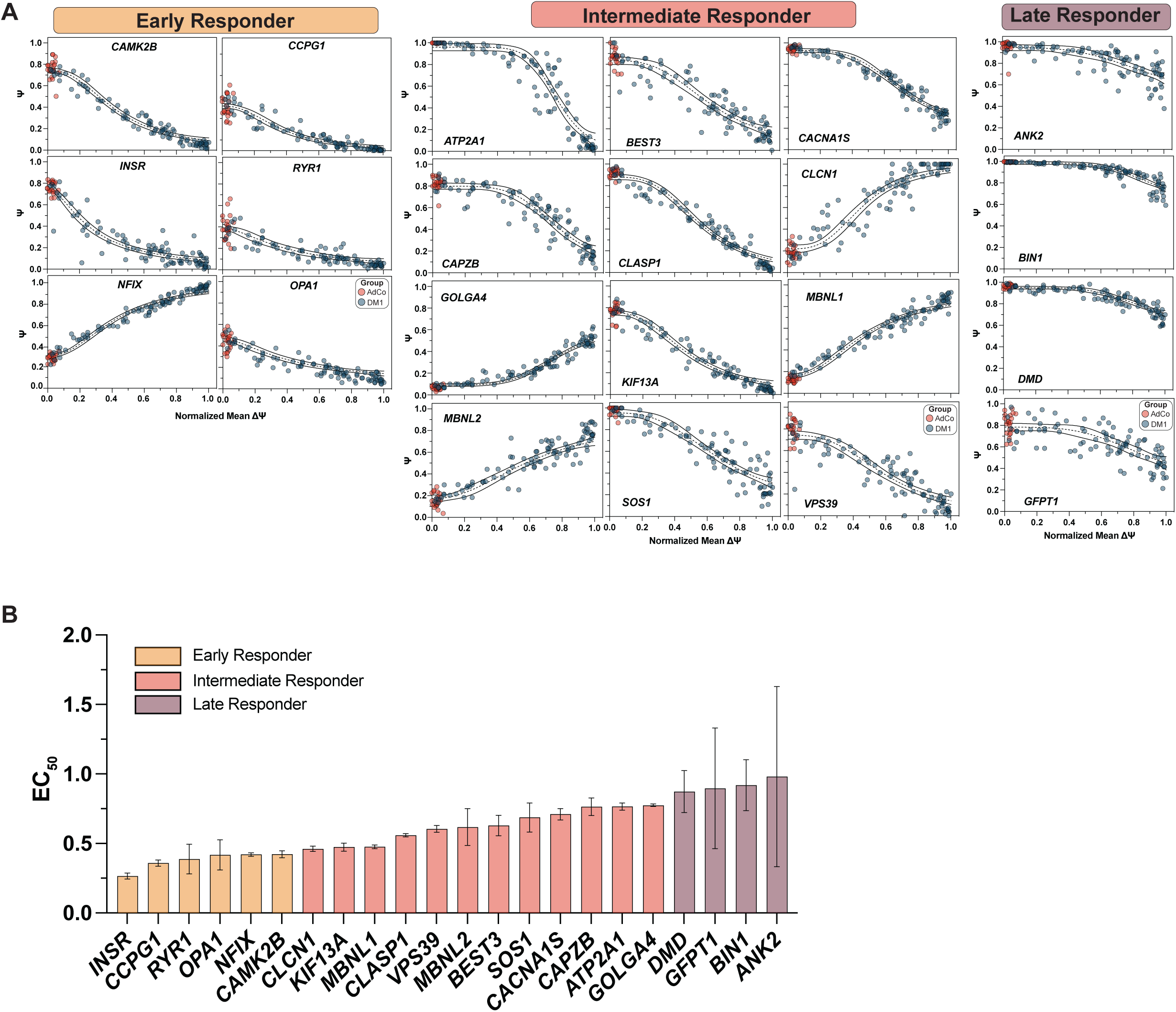
22-events encompassing composite Splicing Index capture RNA splicing shifts across the DM1 disease spectrum. a) Dose response curves of 22 events included in the composite SI. Ψ values derived from total RNAseq (Sup Table 3) are plotted against a normalized mean ΔΨ (Sup Table 1) as a proxy for total transcriptomic dysregulation. Data was fit to a four-parameter dose-response curve to derive curve fitting parameters (ref) (Sup Table 4). Events are classified as early, intermediate, and late responder events based on distribution of EC_50_ relative to the observed median of the full panel. Early responder events have values in lower quartile and late responder events have values in the upper quartile. Curve fit is annotated as the dashed line and the 95% CI of the fit is shown as the solid lines. b) Bar plot of EC_50_ values reported as mean ± SEM. Bar plots are colored based on event response classification. A similar plot of each event’s slope is reported in Sup Fig 3.

Given the known variability of RNA splicing response to marginal changes in functional MBNL concentrations, splice events were prioritized for incorporation within this panel based on the sensitivity of splicing changes across spectrum of disease severity^21,29^. To quantify the responsiveness of each event, Ψ values derived from total RNAseq were plotted against mean ΔΨ as a proxy for total splicing dysregulation; mean ΔΨ has been previously shown to highly correlate with estimated concentrations of functional MBNL in DM1 skeletal muscle (**Fig 2A** & **Sup Table 3**)^21^. Each event was fit to a four-parameter dose-response curve to quantify parameters reflective of biological phenomena, including an EC_50_ value representative of the relative midpoint of the dose response (**Fig 2B** & **Sup Table 4**)^21,22^.

Given our aim to evaluate a biomarker capable of detecting RNA splicing shifts across the full spectrum of DM1 spliceopathy, we used the relative EC_50_ values derived from these analyses to select early, intermediate, and late response events (**Fig 2A**). The group of 22 events encapsulated a complete distribution of EC_50_ values (**Fig 2B**). Most of these splice events were deemed intermediate responders with stepwise changes in Ψ observed across the full disease spectrum, including *CLASP1*, *MBNL2*, and *CAPZB* with EC_50_ values near the median of the complete panel (median EC_50_ = 0.61, 25^th^ percentile = 0.42 and 75^th^ percentile = 0.77). These events accurately convey the largest proportion of splicing dysregulation within the complete DM1 cohort and were the most strongly correlated with global splicing dysregulation (**Sup Table 4**). However, early and late responsive events (i.e. *INSR* and *BIN1*, respectively) were also included to facilitate evaluation of mild and severe DM1 spliceopathy, despite a weaker correlation with total dysregulation. Specifically, *BIN1* is only mis-spliced in the most severely affected DM1 participants (EC_50_ = 0.92) while *INSR* is fully mis-spliced in 90% of this cohort (EC_50_ = 0.27). In total, inclusion of these 22-events variable response events provides a tool to quantify the diverse patterns of mis-splicing observed across the disease spectrum.

### Composite Splicing Index robustly and reliably captures global alternative splicing dysregulation using targeted RNAseq

We next sought to assess if the 22 skipped exon event panel could reliably capture patterns of mis-splicing using targeted, amplicon-based RNA sequencing. To accomplish this objective, we adapted a similar methodology to that previously described in a DM1 mouse model^27^. Multiplex RT-PCR was performed with a low RNA input followed by targeted amplicon RNAseq of event inclusion and exclusion products for all DM1 samples (n = 95) and all available unaffected controls (AdCo, n = 22). As detection bias is a potential disadvantage of this approach due to preferential amplification and sequencing of smaller, exon exclusion isoforms, we compared Ψ values between total and targeted RNAseq datasets. Consistent with this expectation, targeted RNAseq Ψ estimates were generally lower, but not all splice events were impacted (**Sup Fig 4A**). The mean under-detection bias was ∼7% across all events and samples. As expected, detection bias towards skipped exon isoforms was increased for longer cassette exons. The two events with the largest cassette exons (*KIF13A* at 120 nt and *NFIX* at 123 nt) showed the strongest under-detection of exon inclusion at 19% and 15%, respectively. Conversely, almost no under-detection was observed for two of the smallest exons within the panel (*RYR1* at 15 nt and *VPS39* at 33 nt) (**Sup Fig 4B**). Despite this limitation of amplicon sequencing, the greater uniformity of read coverage across relevant splice junctions compared to total RNAseq, reduced confidence limits for Ψ estimates, and significantly higher read counts per splicing event compensate for this issue. Taken together these results support the feasibility of targeted RNAseq as an approach to accurately capture Ψ of select splice events from small inputs of muscle biopsy RNA.

We next employed an overall composite metric to summarize the extent of splicing dysregulation captured by all 22 events, the Myotonic Dystrophy Splicing Index (SI). To derive the SI score for all DM1 participants, Ψ values were normalized to standardized reference values to scale an individual Ψ within boundaries of splicing dysregulation observed between unaffected, healthy controls (Ψ _Median Control_) and the most severely affected DM1 subjects (Ψ _DM95_, 95th percentile within the DM1 sample distribution). The complete spectrum of normalized Ψ values observed within our collective sample group can be visualized via heatmap (**Sup Fig 5A** & **Sup Table 5**). To generate the final SI, normalized Ψ values for each subject were then averaged to generate a linear value from 0 to 1, where values closer to 1 represent those DM1 individuals with the most severe mis-splicing. Normative reference Ψ values for all 22 events were established using all samples subjected to targeted RNAseq (n = 22 AdCo & 95 DM1) (**Sup Table 6**). These reference values were similar to those derived from secondary, non-overlapping AdCo & DM1 cohort (n = 172 DM1 & 25 AdCo, Ψ _DM95_ ICC = 0.996, Ψ _Median Control_ ICC = 0.996, p < 0.0001)^28^. SI values derived using either normative reference set showed high concordance (Pearson *r* = 0.99) (**Sup Fig 5B**).

Consistent with the range of splicing dysregulation observed via total RNAseq, SI scores for DM1 subjects spanned the full range (SI = 0.0 – 1.0) while all AdCo samples exhibited consistently low SI scores indicative of the absence of splicing dysregulation (SI = 0.0 – 0.078) (**Sup Table 1**). A high degree of test-retest reliability was observed for technical replication of targeted RNAseq library preparation, sequencing, and SI calculation (ICC = 0.999, p < 0.0001, n = 6 DM1 and 1 AdCo) (**Sup Fig 5C**). As this metric’s primary objective is to accurately capture the spectrum of splicing dysregulation in DM1 individuals, SI scores strongly correlated with mean ι1Ψ (Pearson *r* = 0.98) (**Sup Fig 5D**)^21,30^. Furthermore, principial component analysis of total RNAseq of 946 significantly mis-spliced events determined that the 89% of the sample variance was defined by the first principal component. Targeted sequencing of the SI component events recapitulated nearly all between participant splicing variance (R^2^ = 0.96) (**Sup Fig 1B** & **Sup Fig 5E-5F**). In total, these results highlight the robust reliability of targeted RNAseq pipeline to accurately determine Ψ values of the 22 splicing events encompassing the SI. Collectively, our findings affirm that this metric serves as a robust, reliable, and disease-specific representation of splicing dysregulation in DM1.

### Myotonic Dystrophy Splicing Index strongly correlates with measures of muscle strength and motor function

To examine the ability of the SI to serve as a biomarker of phenotypic disease severity we next assessed correlations to skeletal muscle performance in all DM1 participants with timepoint-matched functional outcome assessments. As DM1 predominantly affects distal muscles, we first evaluated strength in hand grip and ankle dorsiflexion. Increased SI values were strongly correlated with weaker ankle dorsiflexion (ADF) and hand grip strength (HGS) (Pearson *r* = -0.719 and -0.716, respectively) (**Fig 3A-3B**). Knee extension (KE) showed a weaker correlation with SI, potentially reflecting a disparity between the strength of the proximal muscle groups assessed via this outcome and mis-splicing in the more distal TA (Pearson *r* = -0.392) (**Sup Fig 6A**). To evaluate motor function as measured by ambulation, we evaluated associations with the 10-meter run/fast walk speed (10MRW). A robust correlation was observed (Pearson *r* = -0.680) (**Fig 3C**). In total these analyses indicate that the degree of mis-splicing as measured by the SI correlates with both physical strength and function.

**Figure 3:**
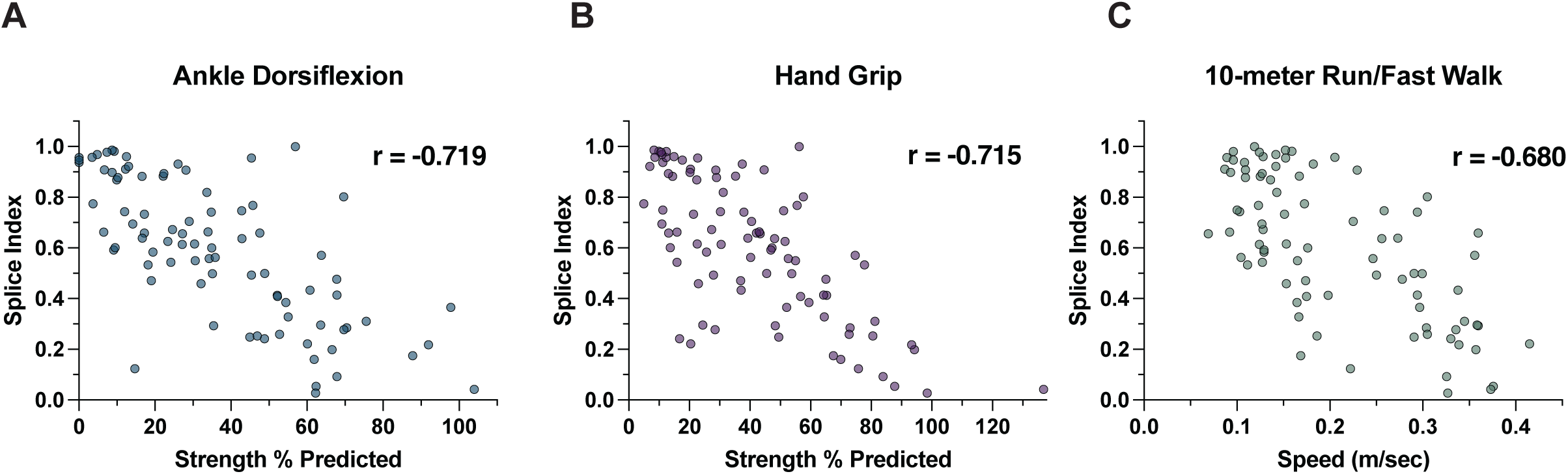
Splicing Index correlates strongly with clinical outcome measures of muscle strength and motor function in cross-sectional DM1 cohort. a-b) Correlation plot of SI versus quantitative measures of ankle dorsiflexion (ADF) and hand grip strength (HGS). Individual measures are reported as the percent of predicted strength as compared to unaffected individuals. ADF Pearson *r* = -0.719 [-0.808, -0.597], p < 0.0001, n = 82 & HGS Pearson *r* = -0.716 [-0.805, - 0.595], p < 0.00001, n = 87. c) Correlation plot of SI versus 10-meter run/fast walk. Individual measures are reported as speed (meters/second). Pearson *r* = -0.680 [-0.782, -0.543], p < 0.0001, n = 82. All correlations are reported as Pearson *r* [95% CI].

We also explored the association between myotonia and the SI. Multiple pre-clinical studies in DM1 mouse models have shown that myotonia rapidly and sensitively responds to administration of disease-modifying therapies, making clinical measures of myotonia and their association with a molecular biomarker ideal for assessment of target engagement in forthcoming clinical trials^23,24^. We found a moderate correlation with a measure of myotonia, the video hand opening time (vHOT) (vHOT_Middle Finger_ Spearman *r* = 0.454 & vHOT_Thumb_ Spearman *r* = 0.377) (**Sup Fig 6B-6C**). As expected, individuals with a high SI had extended times to open a closed fist and individuals with minimal splicing dysregulation displayed low to no clinical myotonia as assessed via this measure. The reduced strength of these associations with the SI, compared to other measures of strength or function, are more likely related to both the clinical variability of myotonia across the spectrum of disease severity and the measurement imprecision of this assessment rather than the lack of true biological association with the underlying pathophysiology. Overall, we observed significant cross-sectional correlations of the SI with multiple functional outcome measures. These associations underscore the relationship between splicing dysregulation in DM1 skeletal muscle and multiple assessments of disease severity commonly used to clinically evaluate DM1 individuals.

While the complete composite SI displayed robust associations with phenotypic performance, we next tested the agreement of individual splice events encompassed within the SI via multiple correlation analysis. Independent of outcome measures, all splice events individually displayed comparable or mildly reduced correlative power compared to the full composite SI. Nearly all events possessed an absolute Pearson correlation coefficient greater than 0.5 for the most strongly correlated outcomes (ADF, HGS, & 10MRW). Regardless of these relatively strong associations, distinct pattens surfaced in this analysis when individual event correlations were ranked by Pearson correlation. Mis-spliced ion channels associated with defects in skeletal muscle excitation-contraction coupling and myotonia– *CANCA1S* e29 & *CLCN1* e7a – consistently emerged either independently or together within the top three most significantly correlated events for ADF, HGS, and 10MWR^18,20^. These findings are consistent with the classification of these events as intermediate responders that show stepwise decrements in Ψ across the full disease spectrum which enhances the overall strength of its correlative power with the majority of DM1 subjects’ performance. In contrast, events classified as early and late responder events, including *INSR* e11 and *BIN1* e11, displayed the weakest independent correlations (**Sup Table 7**).

### SI sensitively captures longitudinal RNA splicing changes in DM1 skeletal muscle over 3-months

We next sought to assess longitudinal stability of the SI by leveraging the inclusion of 35 DM1 individuals in the assembled cohort that provided longitudinal biopsies at baseline (BL) and 3-month (3M) follow up visits (Longitudinal sub-cohort, **Fig 1A**). As a slowly progressive disorder with minimal declines in muscle strength and motor function over 1 year, we predicted that no significant changes in phenotypic presentation would be detected over this short time course^39–41^. In concordance with this hypothesis, no significant changes were observed for all tested clinical outcome measures and all assessments showed high test-retest reliability between BL and 3M (**Fig 4A** & **Sup Fig 7**). SI scores strongly correlated with timepoint matched functional assessments at a comparable strength to the full cross-sectional cohort despite the reduced sample size (**Fig 4B** & **Sup Fig 7**).

**Figure 4:**
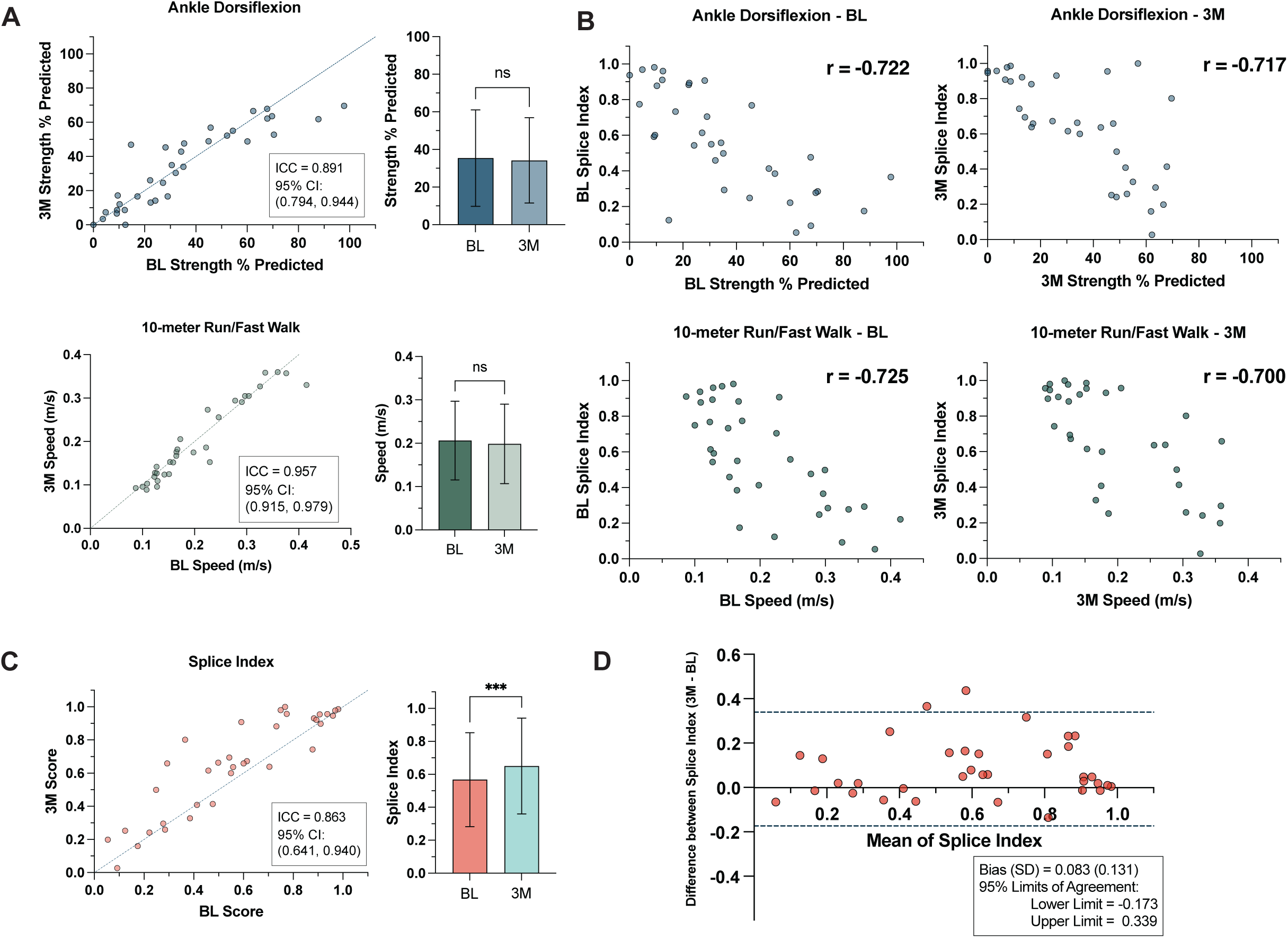
Mean Splicing Index increases between baseline and 3-months in longitudinal sub-cohort with no changes in functional endpoints. a) Correlation plot of baseline (BL) versus 3-month ADF & and 10MRW outcome measures in longitudinally sampled DM1 cohort. Line of agreement (x = y) and ICC with 95% CI are displayed. Bar plot of mean outcomes at BL and 3M is also shown for each outcome measure. Data presented as mean ± SD, ADF n = 34, p = 0.538 & 10MRW n = 32, p = 0.233; paired t-test, ns = not significant b) Correlation plot of BL SI and 3M SI versus timepoint matched measures of ADF and 10MRW. Individual measures are reported as the percent of predicted strength as compared to unaffected individuals (ADF) or as speed (m/s). BL SI v. BL ADF Pearson *r* = -0.722 [-0.852, -0.507], p < 0.0001, n = 34 & 3M SI v. 3M ADF Pearson *r* = -0.717 [-0.849, -0.499], p < 0.0001, n = 34. BL SI v. BL 10MRW Pearson *r* = -0.725 [-0.854, - 0.513], p < 0.0001, n = 34 & 3M SI v. 3M 10MRW Pearson *r* = -0.700 [-0.843 -0.464], p<0.0001, n = 34. All correlations are reported as Pearson r [95% CI]. c) Correlation plot of BL SI versus 3M SI. Line of agreement (x = y) is shown for visualization of SI shift. Test-retest reliability is reported as ICC with 95% CI, n = 35. Bar plot of mean SI at BL and 3M shows significant increase in SI score. Data presented as mean ± SD, n = 35, p = 0.0007; paired t-test, ***p < 0.001 d) Bland Altman plot illustrating agreement between SI scores at BL and 3M in longitudinal sub-cohort.

**Table 1:** Linear Regression Models for 3-Month ADF Measures. Results from three linear regression models examining predictors of 3-month ADF (Average % Predicted) measures (Fig 7). The outcome variables and their corresponding models, parameter estimates, and goodness-of-fit metrics are detailed below. Model 1: The outcome variable for Model 1 is the 3-month ADF Strength % Predicted. The regression equation is defined as 3M ADF (Average % Predicted) = Intercept + BL Splice Index. Model 2: The outcome variable for Model 2 is also the 3-month ADF Strength % Predicted. The regression equation is 3M ADF (Average % Predicted) = Intercept + BL Splice Index + BL ADF Strength % Predicted. Model 3: The outcome variable for Model 3 is the delta (3M-BL) ADF Strength % Predicted. The regression equation is ΔADF = Intercept + BL Splice Index + BL ADF Strength % Predicted. Parameter estimates include Intercept (β0), BL Splice Index (β1), BL ADF Strength % Predicted (β2). Multiple statistical elements are reported (standard errors (SE), 95% confidence intervals (CI), t-values, and P values). An extended intercept table is provided in Sup Table 9.

|  |  |  |  |  |  |  |  |  |
| --- | --- | --- | --- | --- | --- | --- | --- | --- |
| Outcome variable |  | 3M ADF Strength % Predicted |  |  |  |  |  |  |
| Model 1 |  | 3M ADF (Average % Predicted) ~ Intercept + BL Splice Index |  |  |  |  |  |  |
| Parameter estimates |  | Variable | Estimate | Standard error | 95% CI (asymptotic) | \|t\| | P value | P value summary |
| $\beta_0$ | | Intercept | 68.85 | 5.502 | 57.64 to 80.06 | 12.51 | 7.208E-14 | **** |
| $\beta_1$ | | BL Splice Index | -61.56 | 8.741 | -79.37 to -43.76 | 7.043 | 5.51E-08 | **** |
| Outcome variable |  | 3M ADF Strength % Predicted |  |  |  |  |  |  |
| Model 2 |  | 3M ADF (Average % Predicted) ~ Intercept + BL Splice Index + BL ADF (Average % Predicted) |  |  |  |  |  |  |
| Parameter estimates |  | Variable | Estimate | Standard error | 95% CI (asymptotic) | \|t\| | P value | P value summary |
| $\beta_0$ | | Intercept | 24.75 | 7.472 | 9.510 to 39.99 | 3.312 | 0.002359 | ** |
| $\beta_1$ | | BL Splice Index | -21.9 | 8.184 | -38.59 to -5.205 | 2.675 | 0.01181 | * |
| $\beta_2$ | | BL ADF Strength % Predicted | 0.6149 | 0.09153 | 0.4282 to 0.8016 | 6.718 | 1.62E-07 | **** |
| Outcome Variable |  | Delta (3M-BL) ADF Strength % Predicted |  |  |  |  |  |  |
| Model 3 | | ΔADF ~ Intercept + BL Splice Index + BL ADF (Average % Predicted) | | | | | | |
| Parameter estimates |  | Variable | Estimate | Standard error | 95% CI (asymptotic) | \|t\| | P value | P value summary |
| $\beta_0$ | | Intercept | 24.75 | 7.472 | 9.510 to 39.99 | 3.312 | 0.002359 | ** |
| $\beta_1$ | | BL Splice Index | -21.9 | 8.184 | -38.59 to -5.205 | 2.675 | 0.01181 | * |
| $\beta_2$ | | BL ADF Strength % Predicted | -0.3851 | 0.09153 | -0.5718 to -0.1984 | 4.207 | 0.0002048 | *** |

While functional endpoints did not demonstrate a statistically significant difference between baseline and 3-months, a significant difference in the mean SI was observed between BL and 3M biopsies indicative of small increase in overall splicing dysregulation within our longitudinal sub-cohort (**Fig 4C**). We initially inferred that this may be the consequence of biopsy type (14-gauge needle aspirate vs. Bergstrom) or leg selection (ipsilateral vs. contralateral) used for repeat biopsies. However, correlation analysis did not suggest the influence of either variable related to muscle biopsy collection on the SI (**Sup Fig 8**).

Consistent with the observed change in SI being representative of a true shift in splicing dysregulation within this cohort, test-retest reliability between BL and 3M SI scores was reduced (ICC = 0.863, p < 0.0001) compared to technical replicates of the same samples (**Fig 4C** & **Sup Fig 5C**). While this ICC is relatively robust, this analysis indicates that >10% of the variance in SI can be attributed to RNA splicing changes in the skeletal muscle occurring in the 3-month span between biopsies. Moreover, within the cohort we found that individuals with SI scores > 0.8 displayed minimal longitudinal variation (**Fig 4C-4D**). As these individuals already possess severely dysregulated splicing at the baseline assessment, it appears that there is limited capacity for exon inclusion to respond to further changes in intracellular [free MBNL] or CUG repeat load.

Consequently, there appears to be a biological ceiling effect such that minimal variation in the SI score is observed at the upper end of the SI score range. In accordance with this hypothesis, exclusion of these severely affected subjects significantly reduces the between-biopsy agreement (ICC = 0.776, p < 0.0001) (**Sup Fig 9**), indicating general asymmetric variation of splicing dynamics over the 3-month period across the spectrum of disease severity.

### Splicing Index stratifies DM1 individuals across the spectrum of disease severity

Given the mean increase in SI observed over 3-months, we sought to define sub-groups of DM1 individuals according to their basal SI. Rather than rely on visual inspection of the SI distribution to define potential sub-cohorts, we applied two related but distinct data-driven approaches to identify empirical phenotype clusters that could then be evaluated for clinical validity. First, we conducted a latent class analysis (LCA) using Ψ values from five SI events (*CAMK2B*, *CCPG1*, *DMD*, *INSR*, and *GOLGA4*) that displayed modest correlations with the composite measure. The LCA was run using either the entire cohort with available targeted RNAseq data or those DM1 participants with matched baseline outcome measures (sensitivity analysis, n = 52). Overall, the fit indices suggested a three-class solution that broadly corresponded to low, moderate, and high global SI values. The class compositions did not significantly differ with respect to biological sex (p = 0.21) or age (p = 0.24), but statistically significant differences in concurrent measures of function were observed (three class solution, BL only) (**Sup Fig 10** & **Sup Table 8**). Next, k-means fold clustering was applied. This analysis also identified 3 sub-groups consistent with the three classes determined by LCA. This agreement between the two methods reinforces the robustness of the sub-group classification [Group 1 SI mean = 0.21, span = 0.01-0.39; Group 2 SI mean = 0.60, span = 0.41-0.75; Group 3 SI mean = 0.92, span = 0.77 – 1.0]. In combination, these analyses revealed three defined SI stratified sub-cohorts: SI_Mild_ = 0 – 0.4, SI_Moderate_ = 0.41-0.75, and SI_Severe_ = 0.76 -1.0. The bounds of these groups are consistent with visualized inflection points observed in the complete cross-sectional cohort at SI scores between 0.35 - 0.45 and 0.7 - 0.8 when evaluating associations with functional outcomes (**Fig 3**).

Using these defined categories, all 35 longitudinally sampled DM1 individuals were sorted into groups based on their associated baseline SI. Participants were evenly distributed into Mild (BL SI_MILD_ n = 11), Moderate (BL SI_Moderate_ n = 13), and Severe (BL SI_Severe_ n = 11). Consistent with the full cohort, no significant differences in mean outcome measures were observed between BL and 3M across all three groups except for a modest change in HGS for SI_Moderate_ (**Sup Fig 11**). However, individuals with BL SI_Moderate_ scores were the only sub-cohort to have a significant increase in splicing dysregulation at 3-months (**Fig 5A**). While a mean change in SI was observed within the SI_Mild_ group, these changes were not statistically significant in part due to the wide distributional shift of SI values observed from BL to 3M as visualized by Kernel density plot within this sub-cohort. As expected, participants classified as SI_Severe_ showed minimal changes in SI as the RNA mis-splicing ceiling is reached (**Fig 5A-5B**). These differences are further emphasized via visualization of individual subject’s directionality and magnitude of SI change. Subjects within SI_Moderate_ show relatively uniform increases in SI between BL and 3M while individuals within the SI_Mild_ sub-cohort present with variable directionality and magnitude of SI change (**Fig 5A**, connected scatter plots). Overall, these results suggest that DM splicing index can sensitively detect longitudinal changes in SI within participants, and that these changes appear to be more pronounced in moderately affected individuals as defined by their degree of mis-splicing.

**Figure 5:**
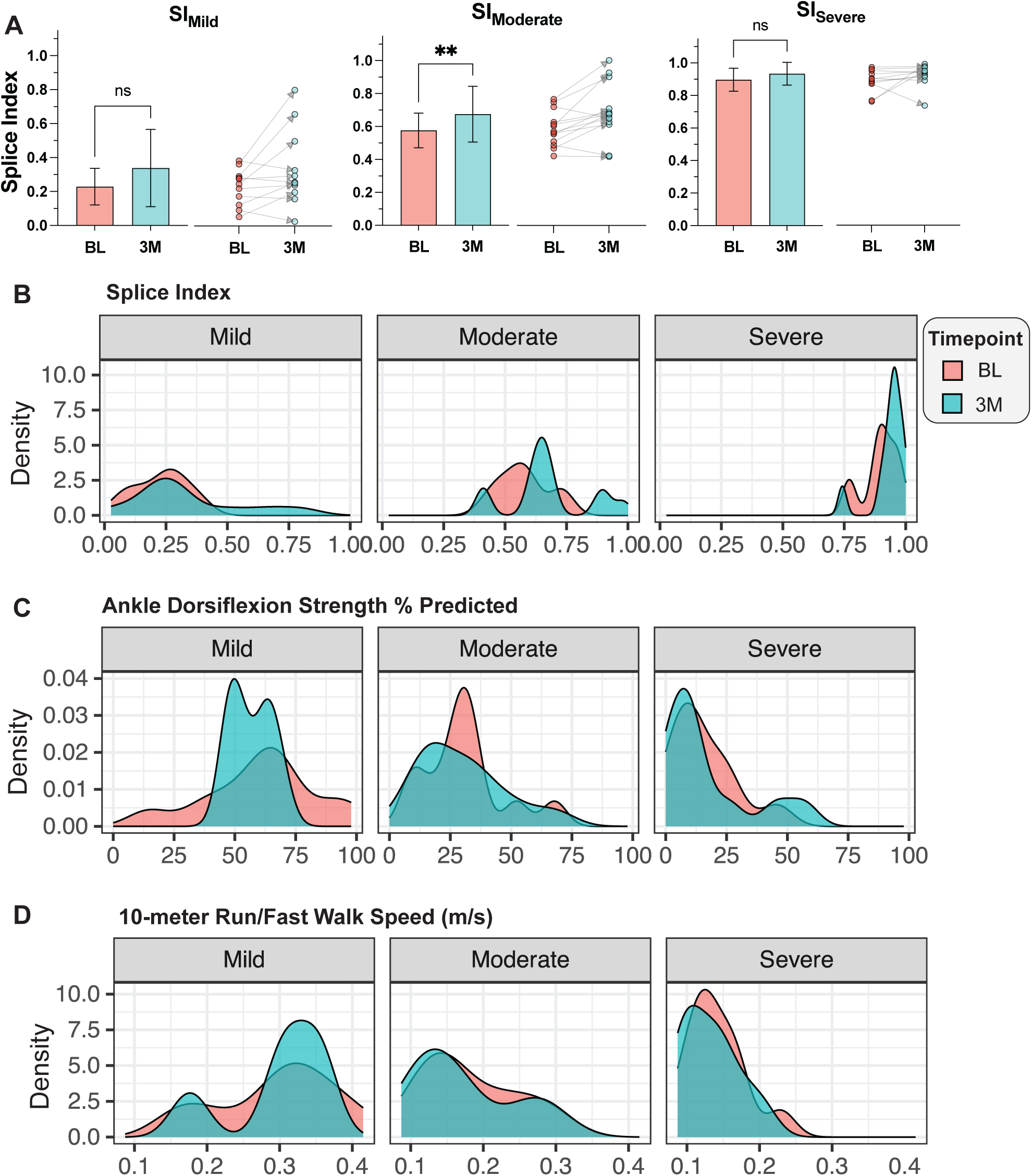
Stratification of longitudinal DM1 cohort by baseline Splicing Index score reveals dynamic changes in splicing dysregulation in mild and moderately affected DM1 individuals. a) Bar plot of mean SI at baseline (BL) and 3-months (3M) for longitudinally sampled DM1 participants stratified by BL SI score: SI_MILD_ (BL SI = 0 – 0.4, n = 11), SI_Moderate_ (BL SI = 0.41 – 0.75, n = 13, and SI_Severe_ (BL SI = 0.76 – 1.0, n = 11). Data represented as mean ± SD; paired t-test, ** p < 0.01, ns = not significant. Connected scatter plots are also displayed to show shifts in BL and 3M paired SI values for each DM1 individual. b) Kernel density estimation plot of BL and 3M SI distributions in Mild, Moderate, and Severe groups. c) Kernel density estimation plot of BL and 3M ADF and 10MRW distributions in Mild, Moderate, and Severe groups. ADF is reported as the percent of predicted strength as compared to unaffected individuals and 10MRW in speed (m/s).

### Individual RNA mis-splicing events differentially capture 3M mis-splicing dynamics in SI stratified groups

Given the dynamic changes in the SI observed in DM1 individuals over 3-months, we aimed to evaluate the sensitivity of early, intermediate, and late responsive splice events encompassed within the SI to assess longitudinal Ψ shifts in SI Mild, Moderate, and Severe sub-cohorts. Normalized ΔΨ (3M – BL) for all 22 events is displayed using box and whisker plots within these sub-cohorts (**Fig 6**). In general, event response classifications accurately reflected the sensitivity of these splice events to detect Ψ changes in the corresponding range of the disease spectrum. That is, early responder events (*INSR* and *CCPG1*) showed the broadest distribution of changes in SI_Mild_, while late responder splice events (*BIN1* and *ANK2*) were able to sensitivity detect 3-month exon inclusion shifts in the most severely impacted DM1 participants (SI_Severe_). Intermediate events detected subtle change across all SI sub-cohorts with variable effect. In concordance with the overall increase in SI over 3-months in the full longitudinal cohort, normalized ΔΨ generally increased across all events within all baseline SI stratified groups.

**Figure 6:**
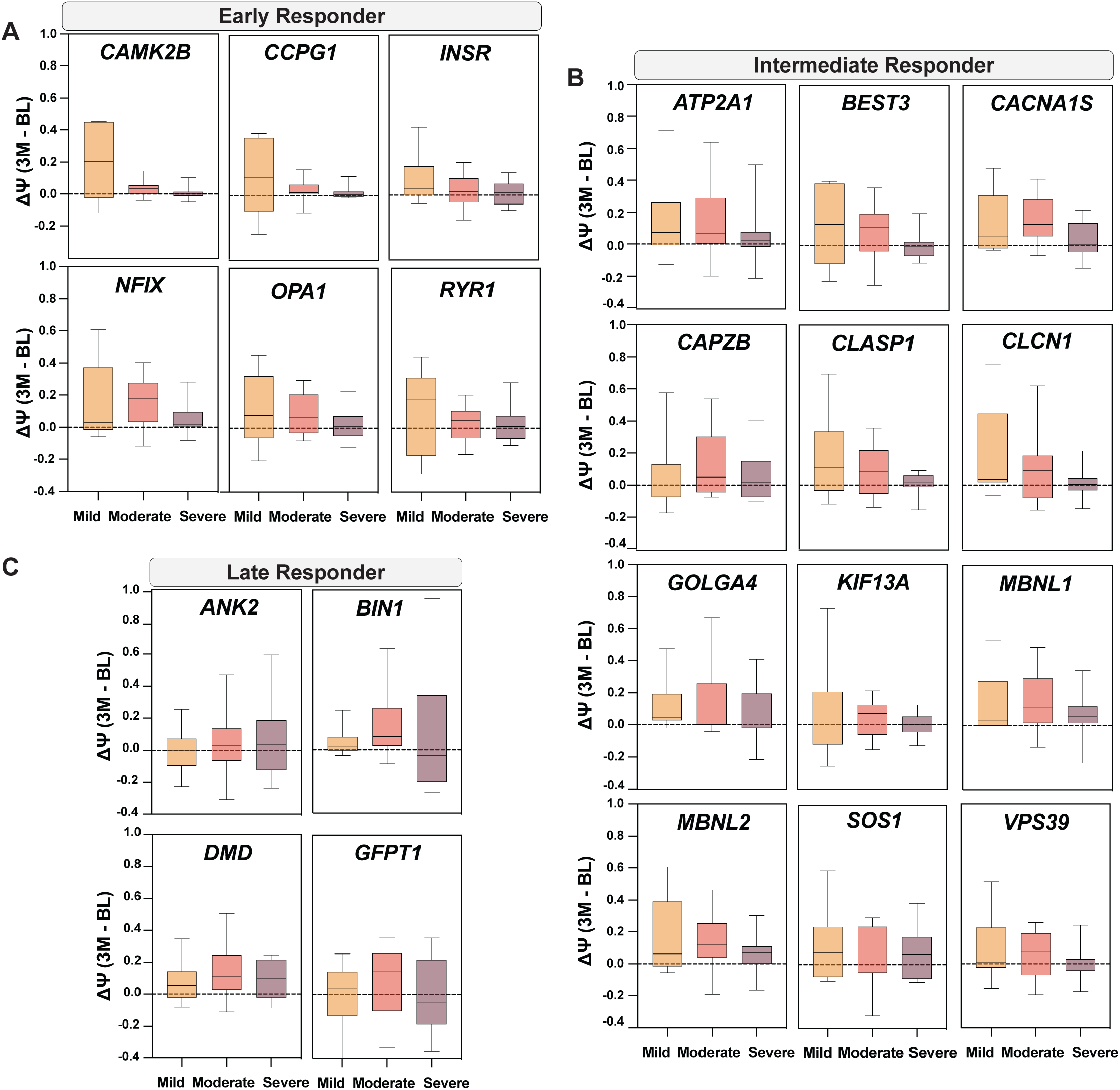
Variable sensitivity of 22 events within Splicing Index panel capture RNA splicing shifts over 3 months in baseline SI stratified sub-cohorts. a-c) Box and whisker plots of normalized ΔΨ (3M-BL) derived from targeted RNAseq (Sup Table 5) for early, intermediate, and late responder events, respectively. Line represents median performance and whiskers extend to max and minimum values. ΔΨ for Mild, Moderate, and Severe SI sub-cohorts are displayed for each event.

However, reversion of mis-splicing was also observed, particularly in SI_Mild_; this observation aligns with the mixed trajectory of SI scores within individuals in this group (**Fig 5A**). Consistent with the significant increase in the composite metric for SI_Moderate_, the magnitude and directionally of normalized ΔΨ was generally uniform. Limited ΔΨ shifts occurred in SI_Severe_ beyond those observed in late responder events in concordance with maximal splicing dysregulation of other events within the index being reached. These observations demonstrate how variably sensitive individual splice events within our composite biomarker may respond to therapeutic target engagement across the spectrum of DM1 spliceopathy using natural disease progression of RNA mis-splicing as a proxy.

### Timepoint comparative analysis of SI indicates that splicing changes precede functional response

Stratification of the longitudinally sampled DM1 cohort by baseline SI revealed that the significant increase in this biomarker over three months was propelled by those individuals with moderate splicing dysregulation (SI_Moderate_). While almost no significant differences in mean outcome measures were observed within the three sub-cohorts (**Sup Fig 11**), an examination of the distribution of performance measures from participants within each SI subsets at both BL and 3M revealed a lack of comparable stratification in the performance range of individuals. This observation was most notable for ADF and 10MRW. Kernel density plots of participant performance on these measures emphasize that the Mild and Moderate sub-cohorts encompass participants with varying measures of muscle strength and motor function spanning the full spectrum of disease severity (**Fig 5C-D**). Consistent with the stability of the SI over 3-months in SI_Severe_, a narrow distribution of outcome measure performance centered at the extreme end of the disease spectrum was observed (**Fig 5C-D**). This is consistent with these individuals reaching a similar ceiling effect for performance on these assessments as that observed for the SI.

These analyses suggest that due to the heterogeneity of disease presentation, especially for less severely affected individuals, splicing dysregulation as measured by SI may be less representative of phenotypic performance in our stratified sub-cohorts at the time of biopsy collection than previously indicated by the full cross-sectional analysis (**Fig 3**). To assess this hypothesis, we leveraged the longitudinally sampled cohort to perform a comparative analysis of both BL and 3M SI scores to timepoint mis-matched outcome assessments. In this analysis BL SI scores exhibited stronger associations for nearly all 3M outcomes – ADF, 10MRW, KE, and vHOT_Middle Finger_ – compared to timepoint-matched measures. In contrast, correlations were diminished when assessing associations between 3M SI scores and BL outcomes (**Sup Fig 12**). Collectively, these analyses suggest that changes in splicing dysregulation precede a phenotypic response, perhaps due to a time-delay between aberrant exon inclusion patterns and downstream biological deficits that modulate phenotypic performance.

### Combination of baseline SI and outcome measures are predictive of 3-month performance

Given the results of our timepoint correlative analysis, we choose to assess the prognostic utility of the SI – i.e. the biomarker’s ability to assess progressive outcomes of affected subjects in the absence of therapeutic intervention. Linear regression modeling was performed using BL SI and ADF, one of the most strongly correlated outcomes observed throughout these analyses. Additionally, we chose to use ADF given its close relationship with strength of the tibialis anterior, the muscle biopsied for derivation of SI in our adult DM1 cohort. Consistent with the timepoint correlation analysis, multiple linear regression modeling using BL SI as a variable indicated the potential power of this biomarker to predict 3M ADF performance (Model 1, adjusted R^2^ = 0.596) (**Fig 7A** & **Table 1**). Inclusion of BL ADF as a covariate significantly increased the power of the model (Model 2, adjusted R^2^ = 0.830) (**Fig 7B** & **Table 1**). While the combination of timepoint-matched SI and ADF strength is successful at predicting future performance at a 3-month assessment, the same was not true when using these variables to predict disease progression as measured by ΔADF (3M – BL) (Model 3, adjusted R^2^ = 0.326) (**Fig 7C** & **Table 1**). This is likely due to the lack of uniform directionality and magnitude of change in SI and skeletal muscle strength as quantified by ADF, especially in the Mild and Moderate sub-cohorts (**Fig 5C**).

**Figure 7:**
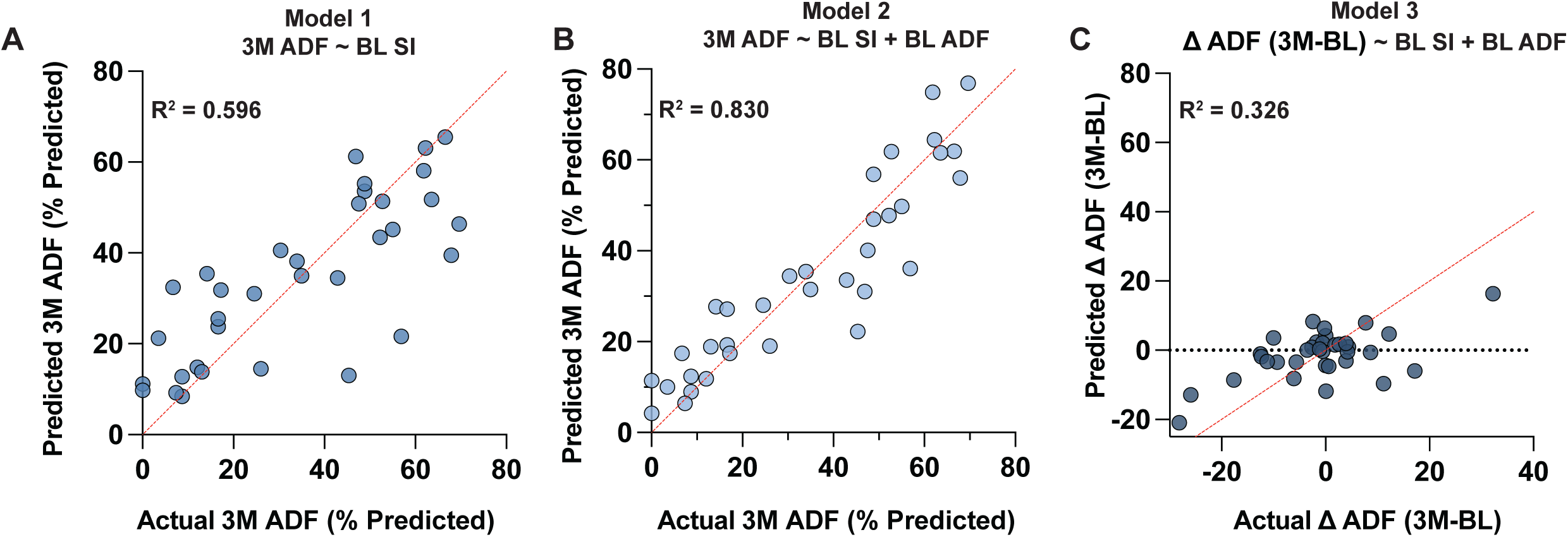
Linear regression modeling highlights prognostic utility of the SI in combination with timepoint matched outcomes to predict future function at 3-months. a-c) Correlation plots of actual versus predicted 3-month (3M) ankle dorsiflexion (ADF) outcomes or progression [ΔADF (3M – BL)] as defined from multiple linear regression Models 1 – 3, respectively. Adjusted R^2^ is reported for all Models. Quantitative parameters of each model are listed in Table 1 and a complete intercept table is provided in Sup Table 9.

## Discussion

These results describe the use and validation of a robust, reliable, and sensitive DM1 composite RNA splicing biomarker – the Myotonic Dystrophy Splicing Index. The DM1 SI accurately distills global RNA mis-splicing severity in skeletal muscle across the disease spectrum and has concurrent validity with measures of muscle strength and function. Early analyses indicate the DM1 SI may be predictive of future function.

RNA splicing is well known to be associated with the pathophysiology of DM1^11^. Prior preclinical work has demonstrated that RNA splicing changes are dynamic in DM1 and each event is differentially sensitive to the degree of MBNL1 sequestration^21,30^. This molecular variability has posed a challenge as to how to capture the entire range of severity by a single splicing-event based biomarker for use in analyzing therapeutic efficacy^29,42^. DM1 phenotypic heterogeneity has also impeded therapeutic progress by causing inflation of study enrollment which has subsequently limited the capability of detecting dose-response relationships and links between changes in RNA splicing and clinical performance. To develop meaningful endpoints in a heterogeneous disease population a biomarker should be able to account for variance at entry and variability in rates of disease progression. These barriers have been overcome by the development of an aggregate measure of RNA splicing inclusive of splice events with variable responsiveness across the disease spectrum. Work described here has demonstrated that the SI is a durable measure of disease-specific splicing dysregulation and is strongly associated with multiple measures of physical function (**Fig 1-3**). We also extend these initial observations by leveraging the inclusion of a longitudinally sampled sub-cohort of DM1 participants to explore the change in SI over a short period of time (**Fig 4-7**).

The SI has high technical replication and reproducibility. This is partly due to high resolution coverage of isoform-specific transcripts containing an exon of interest using targeted RNAseq and subsequent use of a standardized set of normative reference Ψ values. These reference values provide the basis for even scaling of each event’s Ψ across the complete spectrum of splicing dysregulation observed in DM1 and equal weighting in the final, averaged metric of 22 events. This is divergent from methodologies previously used to capture DM1 mis-splicing such as [MBNL]_inferred_, where an exhaustive internal set of DM1 standards representative of the molecular heterogeneity is required in conjunction with a Bayesian inference approach to accurately define an individual subject’s level of RNA mis-splicing^21^. Because the latter analysis is sensitive to appropriate sampling of DM1 subjects across the spectrum if disease for appropriate scaling, changes due to therapeutic engagement may be inappropriately exaggerated or lost, especially when using a small sample size. Normative reference Ψ values described here overcome this barrier by establishing standardized boundaries of RNA mis-splicing for each event from a large, well-phenotyped DM1 cohort like that assembled in this study.

Adults with DM1 report issues associated with mobility, ambulation, or hand function consistently among the top five most impactful symptoms^43,44^. Accordingly, the SI has concurrent validity with measures of muscle strength or ambulation as demonstrated by strong associations with hand grip strength, ankle dorsiflexion strength, and 10-meter run/fast walk (**Fig 3**). Interestingly, there is a modest correlation with a measure of myotonia, the vHOT (**Sup Fig 6**). The rapid correction of this phenotype in DM1 mouse models post therapeutic administration supports the idea that myotonia may provide a rapid assessment of therapeutic target engagement^24,26^. As such, there is interest in applying this outcome measure to human clinical trials^28,45^. However, this outcome measure is variable over time and the diminished strength of association with the SI is likely related to the challenge of quantifying this phenotype in a reliable fashion. Future work aimed at improving reliability of this outcome assessment is needed to fully assess its degree of correlation with the SI. Additionally, as the SI is designed to capture changes association with skeletal muscle performance, associations with other measures of multisystemic dysfunction, such as cognition, would likely be weaker.

While the same principle may apply to develop composite splicing indices indicative of disease severity in other organ systems, the ability to acquire other affected tissues may limit the development of parallel biomarkers outside of skeletal muscle.

One key observation from these analyses is that RNA splicing in DM1 skeletal muscle is more dynamic over short time periods than previously predicted, particularly for a subset of individuals who are moderately affected. The clinical progression of DM1 is measured over years and it is unexpected that a significant difference would be seen over 3-months’ time, which is borne out by clinical observations^39–41^. However, it appears possible that there is a dynamic range in RNA splicing that may portend significant change in function over a 3-month period that appears isolated to subjects with moderate spliceopathy (SI_Moderate_). This is similarly seen, to a lesser extent, in those with a mild spliceopathy (SI_Mild_). Conversely, individuals with a severe spliceopathy do not exhibit significant fluctuations in either the SI or functional outcomes (SI_Severe_) (**Fig 5** & **Sup Fig 7** & **Sup Fig 11**). There are implications for clinical trial design in these findings. Firstly, those individuals within a broad dynamic range (e.g. SI_Moderate_) are likely to be most responsive to disease modifying interventions but also may have the most heterogeneity in clinical trial performance on primary endpoints. This may influence enrollment size in forthcoming therapeutic trials. Additional efforts in a larger DM1 cohort should consider how to best identify these individuals via clinical outcome assessments alone. However, inclusion of early, intermediate, and late responsive splice events within the composite SI show promise in detecting splicing changes across the complete molecular and phenotypic distribution of DM1 disease presentation (**Fig 6**). These splice events may have utility when evaluating therapeutic efficacy in affected individuals with variable disease severity in post-hoc analyses^28,46^.

Timepoint correlation analysis revealed stronger associations between the baseline SI and 3-month functional endpoints than with timepoint matched measures (**Sup Fig 12**). One hypothesis for this difference is that the RNA transcripts measured in this assay are not yet translated to protein, and therefore represent a future composition of muscle function. The time from transcription to translation of the RNA targets may occur over a variable period but are subsequently accounted for in the strength and functional measures at 3-months. This suggests that the SI in combination with baseline functional measures may be predictive of future functional performance, and our regression analyses support this prognostic utility (**Fig 7** & **Table 1**). The ability to predict long term outcomes (e.g., > 12 months) is an open question that requires additional, longer-term studies.

While analyses here validate the utility of the SI for adult-onset DM1, we have preliminarily evaluated the potential validity of the SI to, at minimum, capture disease-specific splicing dysregulation in a small cohort of congenital myotonic dystrophy (CDM) and ambulatory DM2 participants (n = 8 CDM & 4 DM2) (**Sup Fig 13**). Ψ values of the 22 events encompassed within the SI effectively cluster these DM sub-types along the range of splicing dysregulation observed in our DM1 cohort. This analysis also provides an initial indication that the SI can sensitivity assess changes in global mis-splicing in these populations; two CDM children with longitudinal biopsies displayed agreement with previously described shifts in global splicing dysregulation during pediatric development (**Sup Fig 13B**)^30^. Although genotyped and aged-matched normative reference Ψ values would be required for use of the SI in these DM sub-types, this data shows promise that either the DM1 SI or a modified composite splicing biomarker could be developed for these populations.

In summary, the SI is a robust, reliable, and sensitive biomarker that captures dynamic RNA splicing changes, is associated with physical strength & ambulation, and is potentially predictive of future function in adult DM1 individual. These properties indicate that the SI can serve as a biomarker of strength and function during the course of a clinical trial. When used in combination with physical outcomes, the SI can serve as a guide when selecting individuals for clinical trials as those with moderate splicing defects are both the most variable but also the most likely to demonstrate early benefit from disease modifying therapies.

## Methods

### Participants and muscle biopsy collection

Participants with clinically or genetically diagnosed DM1 over the age of 18 were enrolled in END-DM1 (NCT03981575) or a prior pilot study (HELP-DM1). Individuals were required to be ambulatory and have sufficient tibialis anterior muscle bulk for muscle sampling. Individuals were not able to participate if they were on anti-myotonia agents or anti-coagulants, or had a platelet count less than 50,000 or history of a bleeding disorder.

Muscle biopsies from DM1 participants were collected with either a 14-gauge argon Supercore needle or a Bergstrom needle of the tibialis anterior (TA) by previously described techniques^28^. At the 3-month longitudinal assessment select biopsies were conducted on the ipsilateral or contralateral TA. Select unaffected adult control (AdCo) and congenital myotonic dystrophy (CDM) biopsies of the TA, vastus lateralis, and soleus were collected as previously described^30^. Additional AdCo, myotonic dystrophy type 2 (DM2), Duchenne muscular dystrophy (DMD), and limb girdle muscular dystrophy (LGMD) biopsies were collected from the TA of ambulatory subjects with a 14-guage needle as part of an in-house biorepository study. Available demographic information of all available participants is listed in **Sup Table 1**.

### Functional Outcome Measure Assessment and Analysis

Quantitative muscle testing (QMT) of hand grip, ankle dorsiflexion, and knee extension strength were conducted with a fixed quantitative system^28^. Strength measures are reported as the percent of predicted strength as compared to a non-affected individuals of the same age, sex, and height^28^. The 10-meter run/fast walk was conducted as previously described and times were used to calculate fast walk/running speed (meters/second)^47^. Myotonia was assessed using the video hand opening time (vHOT) assessment^28^. In brief, subjects are instructed after resting their dominant hand in an open position for 5 minutes on a flat surface to make a fist and squeeze for 3 seconds. Participants are then cued to open their hand completely until all fingers and thumb are fully extended and relaxed. A blinded rater reviewed video of the assessment and measured time from when the participant is cued to open the hand until the middle finger is fully extended (vHOT_Middle Finger_) and until the thumb is fully extended (vHOT_Thumb_). Following a 5-minute rest a second trial was completed. Time expressed in seconds from the second trial for both vHOT_Middle Finger_ and vHOT_Thumb_ were used for analyses. All measures are reported in **Sup Table 1**.

### Muscle Biopsy Processing

RNA was extracted from biopsies via a TRIzol-chloroform extraction supplemented with a bead-based tissue homogenization (Benchmark Scientific) and purified using the Zymo Clean and Concentrator – 5 kit following the standard protocol for on-column DNase treatment. RNA quality and abundance was assessed using fragment analysis (Agilent Fragment Analyzer 5200, DNF-472 (15 nt) RNA kit) and Qubit 4 Fluorometer (Invitrogen). Most RNA samples had RIN values >7, but samples were utilized for subsequent analyses if they had RIN scores > 5.5. RNA was stored at -80°C until further processing.

### Total RNAseq library preparation, sequencing, and RNA splicing analysis

Libraries were prepared using up to 250 ng of RNA using the NEBNext rRNA Depletion Kit in conjunction with the NEBNext Ultra II Directional RNA Library Prep Kit for Illumina (New England Biolabs). Deviations from the manufacturer’s protocol include modifications of the recommended 5-fold adaptor dilution to a 40-fold adaptor dilution for RNA inputs between 250ng-10 ng. Libraries were paired-end (2 x 75 bp) sequenced on the NextSeq 2000 using a P3 Reagents at a loading concentration of 750 pM with a 1% PhiX spike-in. Samples were sequenced for coverage at > 65 million reads. Library quality assessment and read mapping were performed as previously described Differential splicing analysis for total RNA-seq data was performed using rMATS as previously described^30^. Event Ψ was calculated and deemed significantly mis-spliced when |ΔΨ | ≥ 0.1, FDR ≤ 0.05 between defined groups. Derivation of mean ΔΨ using the top 50 most dysregulated splicing events from this output was calculated as previously described^30^. For splicing event classification, Ψ values derived from total RNAseq (**Sup Table 3**) were plotted against the normalized mean ΔΨ scaled from 0 to 1 and fit to a four-parameter dose response curve as previously described to derive quantitative parameters of the splicing shifts over the disease spectrum (**Sup Table 4**) (ref).

### Targeted RNAseq library preparation and sequencing

Targeted RNAseq library was designed similarly to that previously described for generation of a species-specific splice index in DM1 mouse models^27^. In brief, RNA splice events identified using total RNAseq dataset as having a large, DM1-specific effect size, low Ψ variability in AdCo subjects, and a gene structure amenable to amplicon sequencing were included in the final 22-event panel^28^. Specific splice events that were previously identified as mis-spliced in DM1 patients and were moderately correlated with manual muscle testing the ankle dorsiflexion in DM1 subjects were also included despite smaller effect sizes^29^.

cDNA was synthesized from 10 ng of high-quality RNA using the SMART-Seq v4 Ultra Low Input RNA Kit (Takara). Genome coordinates for alternative exons and PCR primers for amplifying cDNA are listed in in **Sup Table 11**. To normalize a balance of sequence reads across the 22 splice events, especially for low abundance transcripts, the 22 primer sets were distributed across 2 multiplex, first-stage PCR reactions of 14 and 16 cycles (PCR1 A and B, respectively). All PCR1 primers included a 5′ adapter sequence for incorporation of sample barcodes and priming sites for Illumina sequencing in a second-stage PCR (PCR2). Libraries were individually normalized based on concentrations assessed using the Qubit (Qubit 4 Fluorometer (Invitrogen) and average library size as determined by smear analysis (Agilent Fragment Analyzer 5200, DNF-474 HS NGS kit). Libraries were paired-end (2 x 151) sequenced on the iSeq 100 using an i1 Reagent v2 (300-cycle) kit at a loading concentration of 125 pM with a 10% PhiX spike-in.

### Quantification of SI

Amplicon library reads in raw FASTQ format were aligned to a custom reference sequence set of exon inclusion and exclusion isoforms (**Sup Table 11**) using HISAT2 v2.1.0 and filtered reads with the -no-spliced-alignment flags and -no-softclip run parameters. Primary aligned reads spanning splice junctions were subset from the resulting bam files using a combination of Samtools v1.7 and Bash CLI. Isoform-specific reads with unambiguous alignment were counted and further processed in a custom python script to derive Ψ for each splice event. Ψ values were calculated as the fraction of exon inclusion isoform reads relative to all exclusion and inclusion isoform counts. The SI score was then calculated as follows. For each subject *y*, normalized Ψ values were calculated for each splice event *z* as (Ψ*_y,z_* – Ψ _median control,*z*_)/( Ψ _DM95, *z*_ – Ψ _median control, *z*_) where Ψ _median control, *z*_ is the median Ψ for event *z* in unaffected healthy adults, and Ψ _DM95, *z*_ is the 95th percentile for most severely affected DM1 subjects^27^. Normative values used throughout these analyses unless otherwise annotated were derived from our cohort of AdCo samples (n = 22) and all DM1 individuals subjected to targeted RNAseq (n = 95) (**Sup Table 6**). Normalized Ψ values for each subject *y* where then averaged to generate a linear value from 0 to 1.

### Statistical Analyses

Clinical data were extracted from Redcap using Microsoft Excel Version 16.75. Any clinical datapoints that were available but deemed to be questionable by the study evaluators were subsequently reviewed by the study’s Principal Investigator. An ultimate decision regarding inclusion/exclusion was made jointly by the evaluators and P.I. for these specific datapoints.

Univariate correlations and multiple linear regression were performed using GraphPad Prism 10.1.0 and R statistical software (R 4.2.3). Univariate Pearson or Spearman correlation coefficients were calculated depending on the distributional properties of the variable, and significance was reported with a two tailed p-value and 95% confidence interval. Intra-class correlation coefficients were calculated using a two-way fixed effects model for absolute agreement between measures. Multiple linear regression modeling assumed a least squares regression and the adjusted R-squared was used to quantify goodness-of-fit. Assumptions of regression including normality of residuals were assessed for each model. P-values < 0.05 were considered significant.

Latent class analysis (LCA) was performed in the R statistical environment (v4.2.1) using poLCA^48^. First, the correlation structure of all 22 RNAseq Ψ values was examined to quantify the extent of collinearity. This pre-analysis step is standard for LCA because the approach is sensitive to multicollinearity. Based on the correlations, five SI splice events exhibiting only modest correlations were selected (*CAMK2B, CCPG1, DMD, INSR, and GOLGA4*). Next, the mean SI value for each event was calculated using the entire cohort. For each individual, their continuous SI value for each event was dichotomized such that one indicated a value greater than the mean SI for that event and zero indicated less than the mean value. The LCA was run using both the entire cohort with available targeted RNAseq data (n = 129) and those adult onset-DM1 subjects with matched baseline measures only (n = 52) (sensitivity analysis). We chose “any class size smaller than 10% of the total sample size” as a predetermined stopping point as the threshold to stop testing additional classes. Based on this stopping criteria, two-, three-, and four-class models were tested, but only two- and three-class solutions were interpreted. Each model was run 300 times with 5,000 set as the maximum number of iterations of estimation algorithm cycles to increase that the odds of finding the best global maximum of the log-likelihood function. Goodness of fit was assessed using Akaike information criterion (AIC) and Bayesian information criterion (BIC). Predicted class membership was validated using concurrent and future functional measures of strength and mobility to evaluate the potential clinical utility of these data-driven phenotypes. Bounds of three defined SI sub-cohorts were determined using sklearn’s KMeans clustering package. The supervised model grouped samples by Splicing Index score into Mild, Moderate, and Severe categories. Minimum and maximum scores in each class informed the boundaries of their respective groups.

## Data Availability

RNAseq datasets and computer code scripts used for analysis in this study can be found in the following databases and repositories:

- Total and Targeted RNAseq libraries: NCBI SRA PRJNA1079722 (https://www.ncbi.nlm.nih.gov/bioproject/PRJNA1079722)
- Analysis Scripts: https://github.com/Center-for-Inherited-Muscle-Research/splice_index

## Acknowledgments

Thank you to the Myotonic Dystrophy Clinical Research Network (DMCRN) for the use of END-DM1 muscle biopsies and associated clinical data.

| DMCRN Consortium Contributing Sites |  |
| --- | --- |
| Institution Name | Site Staff Name, Degree (Role) |
| University of Rochester Medical Center | Johanna Hamel, MD (Site PI) |
|  | Jeanne Dekdebrun (Primary CRC) |
|  | Katy Eichinger (Primary CE) |
| Virginia Commonwealth University | Nicholas Johnson, MD, M. Sci, FAAN (Site PI) |
|  | Carino Jennings (Primary CRC) |
|  | Aileen Jones (Primary CE) |

## Author Contributions

**Marina Provenzano:** Investigation, project administration, methodology, data curation, formal analysis, visualization, writing; **Kobe Ikegami:** Investigation, data curation, formal analysis, visualization, writing; **Kameron Bates:** Methodology; **Alison Gaynor:** Data curation; **Julia M. Hartman:** Data curation**; Aileen S. Jones:** Data curation; **Amanda Butler:** Data curation; **Kiera N. Berggren:** Data curation; **Man Hung:** Formal Analysis; **Dana M. Lapato:** Formal Analysis, visualization, writing; **Michael Kiefer:** Data curation, methodology; **Charles Thornton:** Methodology, conceptualization, funding acquisition; **Nicholas E. Johnson:** Conceptualization, supervision, project administration, data curation, funding acquisition, writing; **Melissa A. Hale:** Conceptualization, investigation, supervision, project administration, methodology, data curation, formal analysis, visualization, writing

## Figure Legends

**Supplemental Figure 1:**
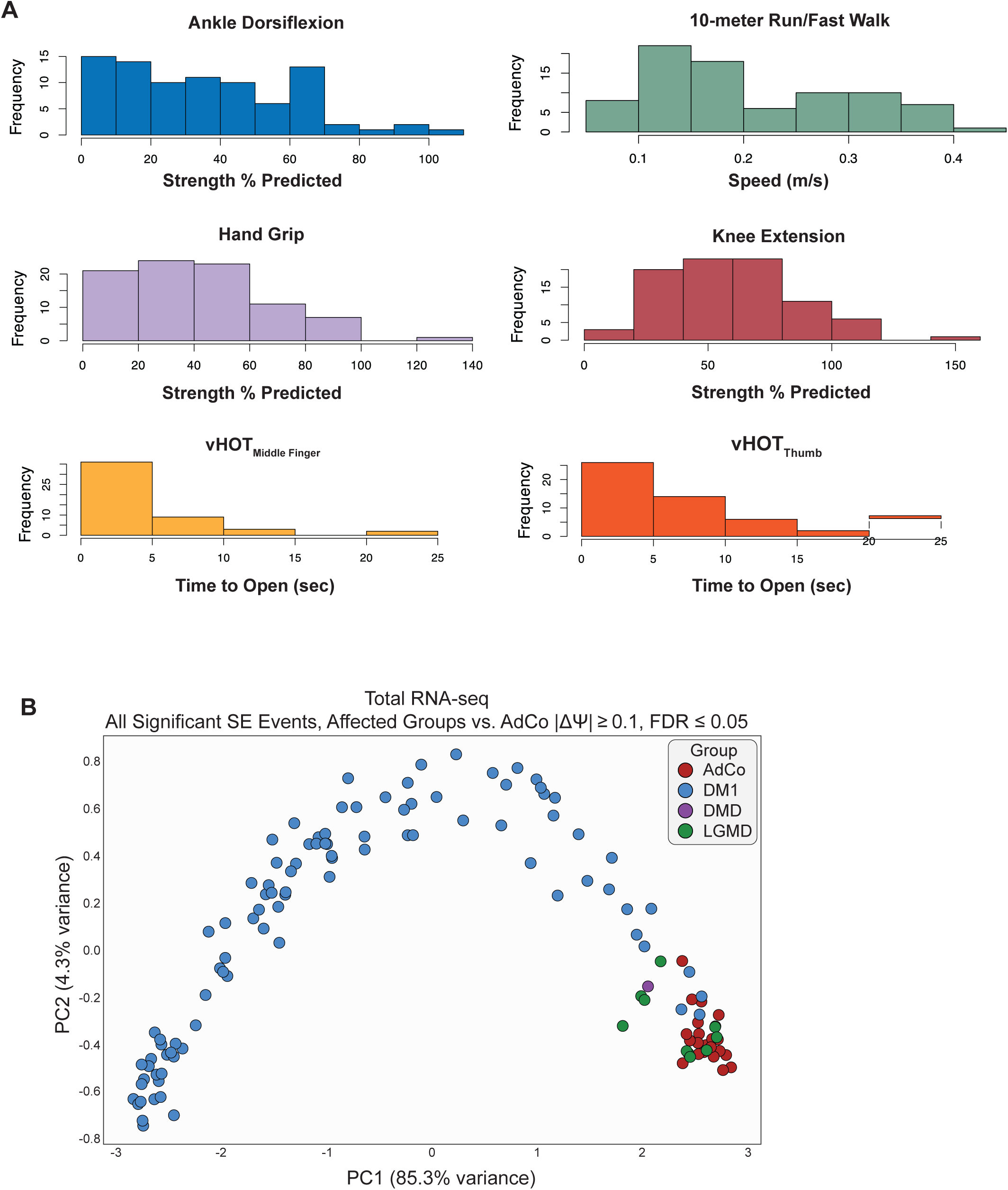
Phenotypic and molecular heterogeneity of DM1 participants subjected to total RNA-seq. a) Distribution of performance measures captured in all DM1 participants via quantitative muscle testing (ankle dorsiflexion, hand grip strength, knee extension), timed motor test (10-meter run/fast walk), and myotonia (video hand opening time – vHOT). b) Principal component analysis of all significantly dysregulated skipped exon (SE) events in muscle biopsies subjected to total RNAseq between DM1 versus unaffected adult controls (AdCo) and disease control reference groups (DMD & LGMD). In total 946 significantly mis-spliced, skipped exon events were defined by |ΔΨ| ≥ 0.1, FDR ≤ 0.05 (Sup Table 2).

**Supplemental Figure 2:**
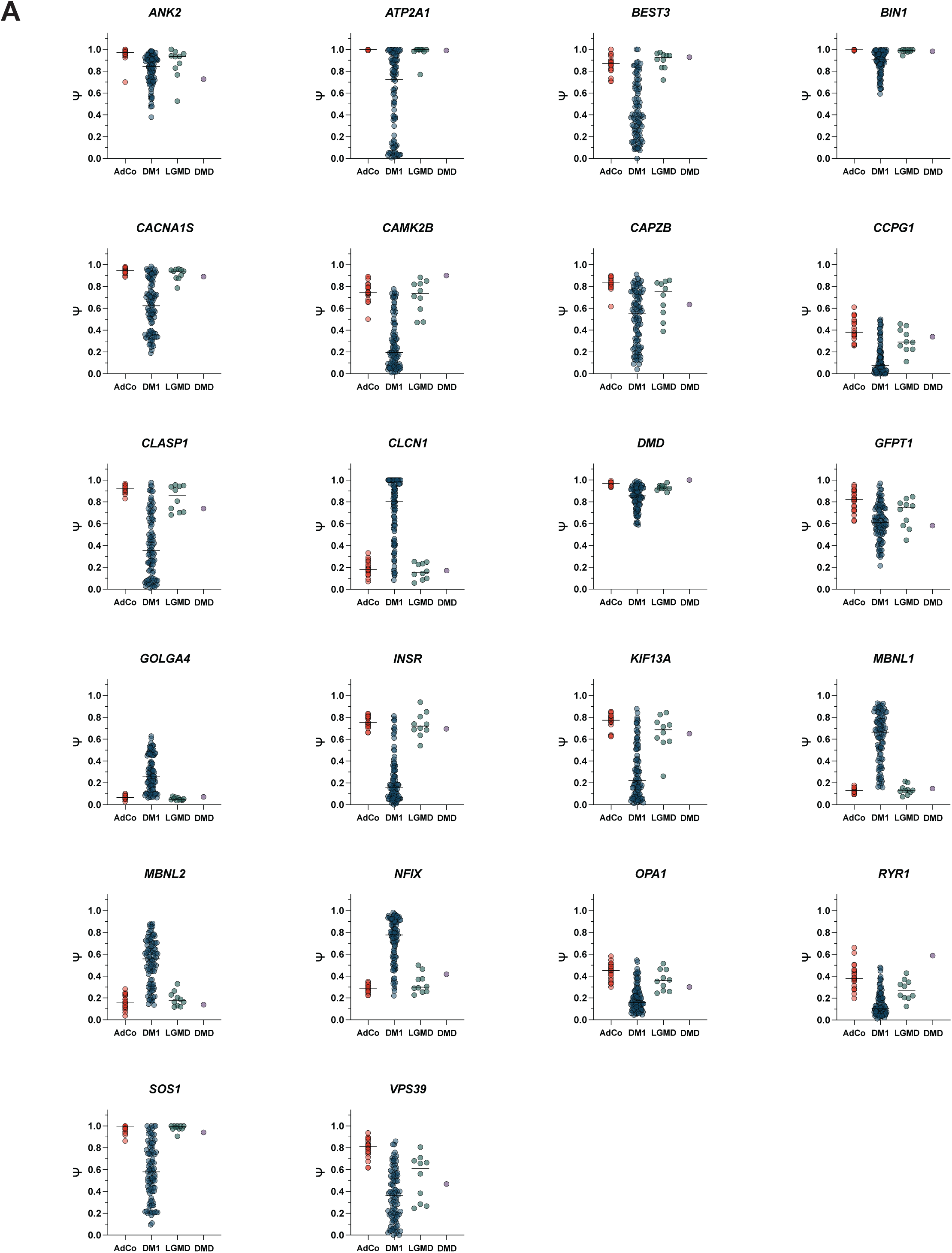
22 splice events selected for composite Splicing Index are disease-specific and display minimal variability in unaffected adult controls and disease controls. Strip plots of Ψ values from total RNAseq for 22 skipped exon events in composite Splicing Index panel. Ψ values (Sup Table 3) are colored by sample group and bar represent median Ψ.

**Supplemental Figure 3:**
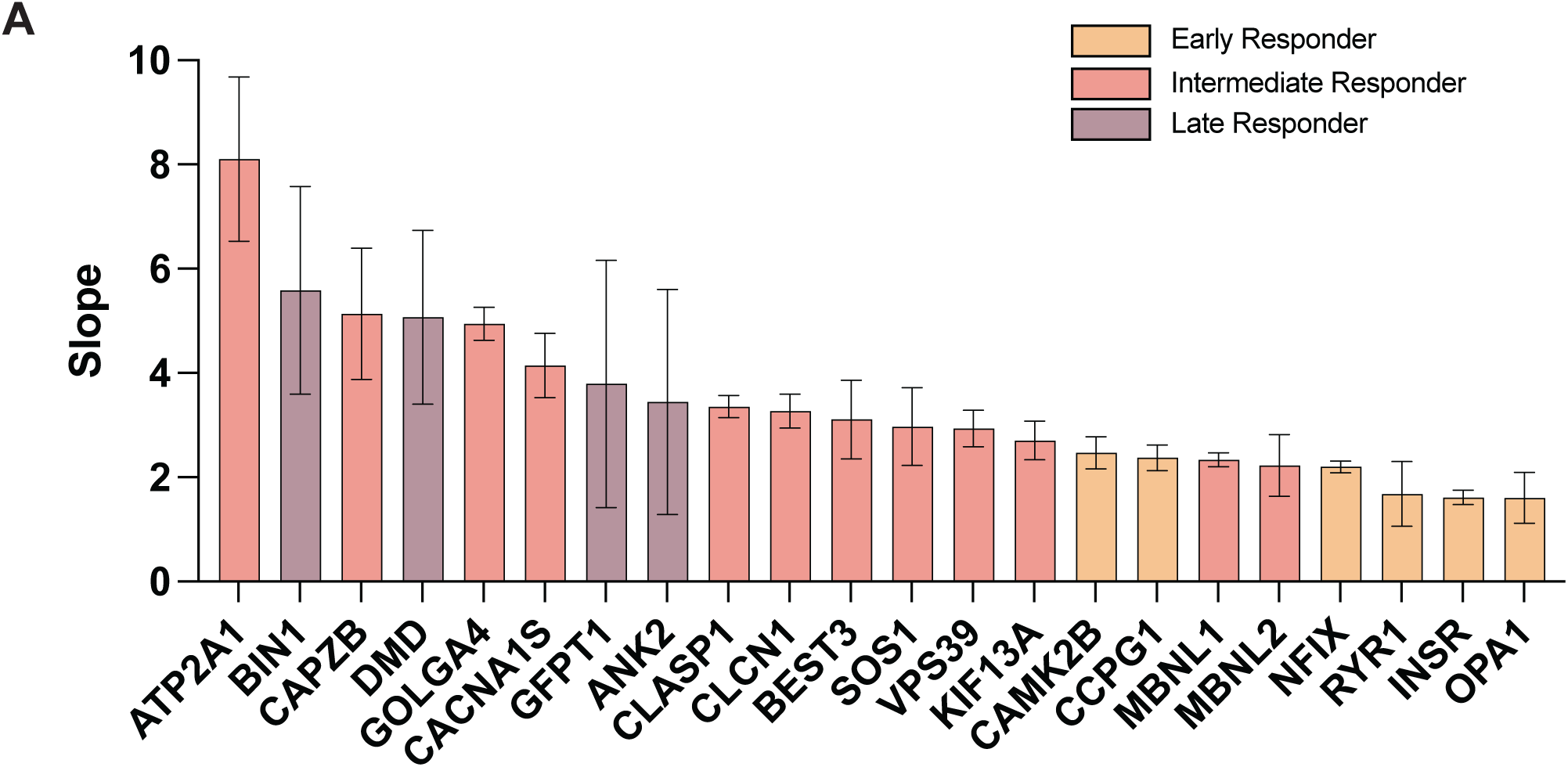
Slope derived from dose response curve fitting of 22 splice events in composite Splicing Index panel. Bar plot of slope values derived from dose response curve fitting and colored based on event response classification (i.e. early, intermediate, and late) (Fig 2A). Data represented as mean ± SEM.

**Supplemental Figure 4:**
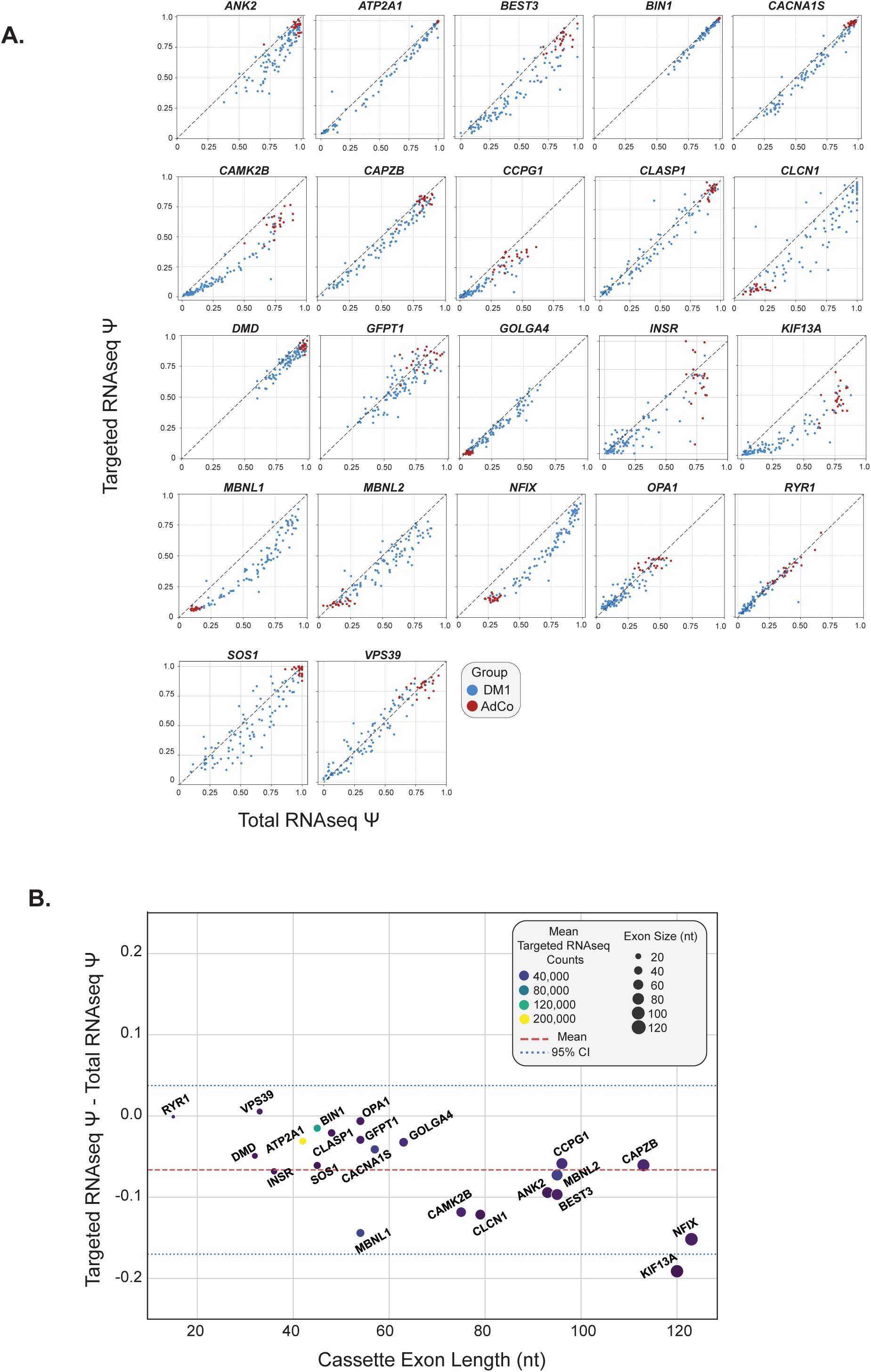
Targeted RNAseq shows limited under-detection bias of exon inclusion isoforms for select splice events included within the Splicing Index. a) Comparison of Ψ estimates derived from total and targeted RNAseq for panel of 22 skipped exon splice events (Sup Table 3 & Sup Table 5). Line of agreement (x = y) is displayed to show relative detection bias of smaller, exon exclusion isoforms in targeted RNAseq and subsequent lower Ψ compared to total RNAseq. b) Targeted RNAseq shows small bias for exon exclusion isoforms that is moderately correlated to cassette exon length. Red dotted line represents mean Ψ difference between methodologies and blue dotted lines represent the mean ± one standard deviation. Size of each event dot is reflective of exon length and color is reflective of mean event counts in targeted RNAseq.

**Supplemental Figure 5:**
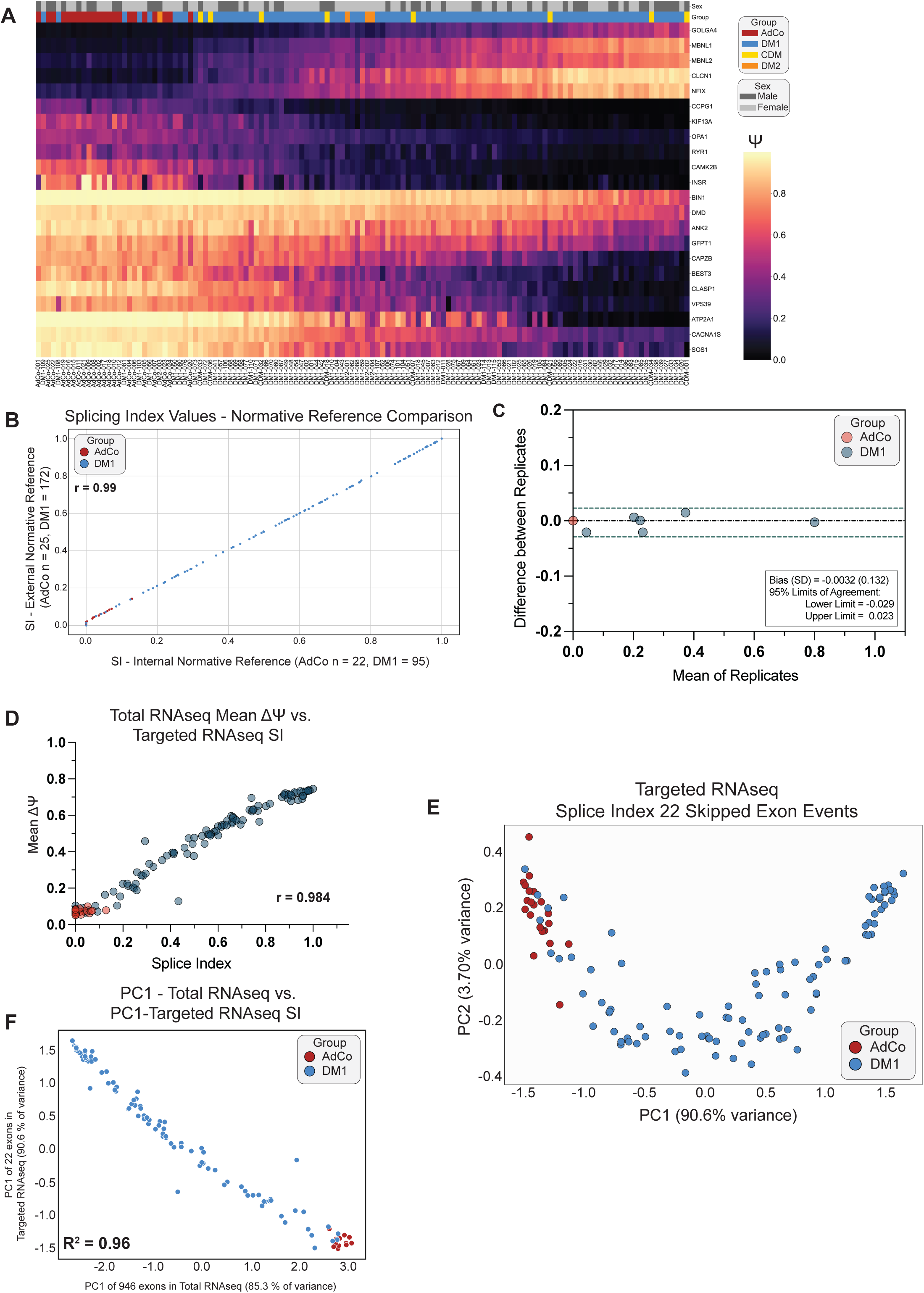
Targeted RNAseq of 22-events captures global spliceopathy in DM skeletal muscle. a) Heatmap displaying normalized Ψ values for 22 SI splicing events derived from targeted RNA-seq of DM1, AdCo, CDM, and DM2 samples subjected to targeted RNAseq (Sup Table 5). Columns (individuals subjects) are force-ranked by SI from 0 to 1 while rows (splice events) were subjected to hierarchical clustering. Sample group and sex are annotated above and Subject ID is listed below. b) Correlation plot of SI for all DM1 and AdCo subjects using internal normative reference set (Sup Table 6) versus scores scaled using external normative reference values (Pearson *r* = 0.99) c) Bland Altman plot illustrating agreement between SI scores derived from technical replications of 6 DM1 and 1 AdCo sample using targeted RNAseq. Inter-rater reliability is reported as an ICC with 95% confidence intervals ICC = 0.99 [0.994, 1.0], p < 0.00001). d) Scatter plot of SI derived via targeted RNA-seq versus mean ΔΨ of top 50 most dysregulated SE events captured from total RNA-seq of the same sample (Pearson *r* = 0.984 [0.978, 0.989], p < 0.0001). e) Principal component analysis of Ψ values for 22 splicing events encompassing the composite Splicing Index derived from targeted RNAseq. f) Correlation plot of principal component 1 from total RNAseq (Sup Fig 1B) versus targeted RNAseq (Sup Figure 5D). R^2^ is reported. All statistical analyses reported with [95% CI].

**Supplemental Figure 6:**
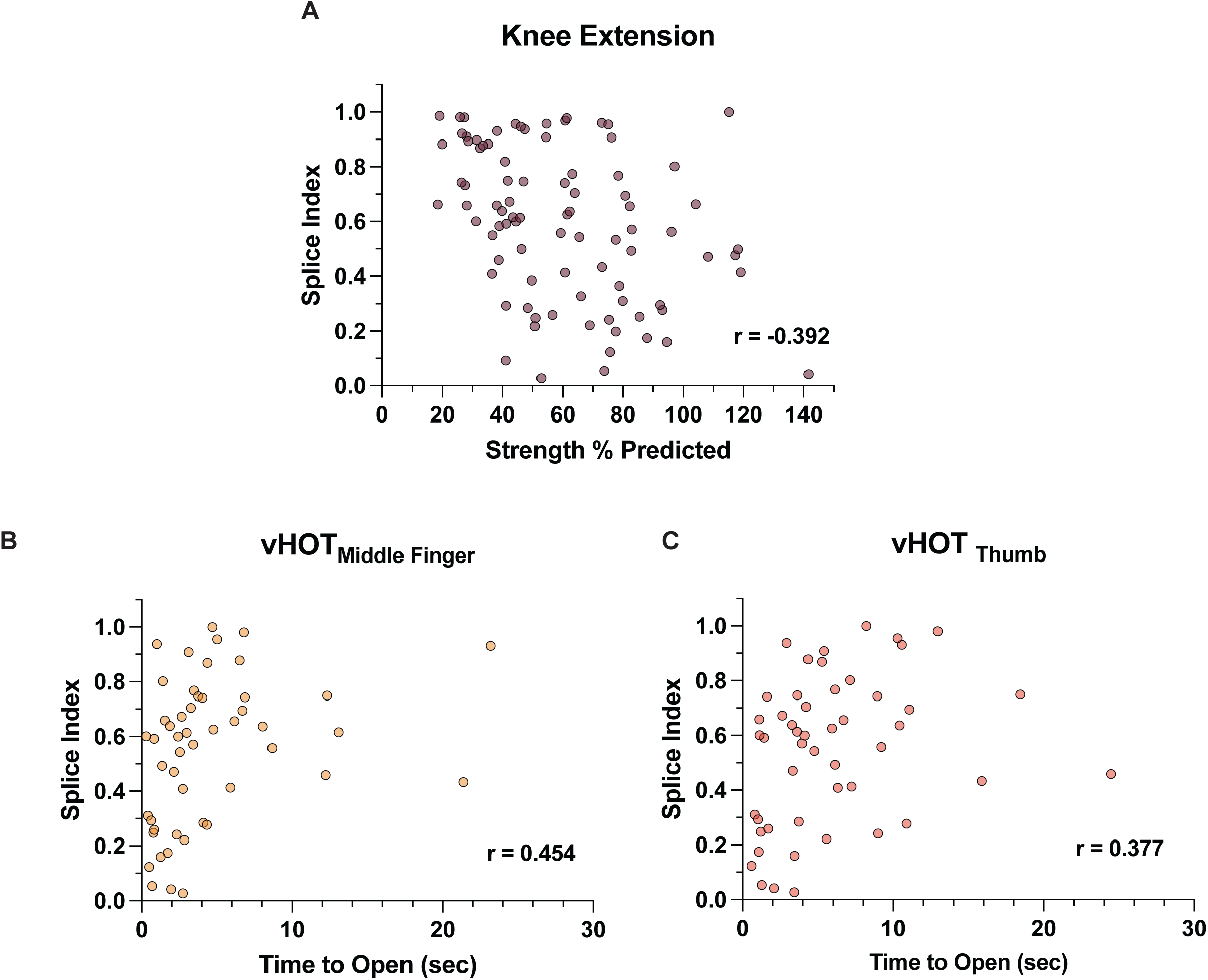
Splicing Index correlates mildly with clinical outcome measures of proximal muscle strength and myotonia in cross-sectional DM1 cohort. a) Correlation plot of SI versus quantitative measure of knee extension (KE) strength. Individual measures are reported as the percent of predicted strength as compared to unaffected individuals. KE Pearson *r* = -0.392 [- 0.556, -0.197], p = 0.0002, n = 87. b-c) Correlation plot of SI versus video hand opening time (vHOT_Middle Finger_ and vHOT_Thumb_). Individual measures are reported as time to open closed fist (seconds). Spearman *r* is reported. (vHOT_Middle Finger_ *r* = 0.454 [0.193, 0.655], p = 0.0009, n = 50; vHOT_Thumb_ *r* = 0.377 [0.099, 0.601] p = 0.0075, n = 49). All correlations are reported as Pearson/Spearman *r* [95% CI].

**Supplemental Figure 7:**
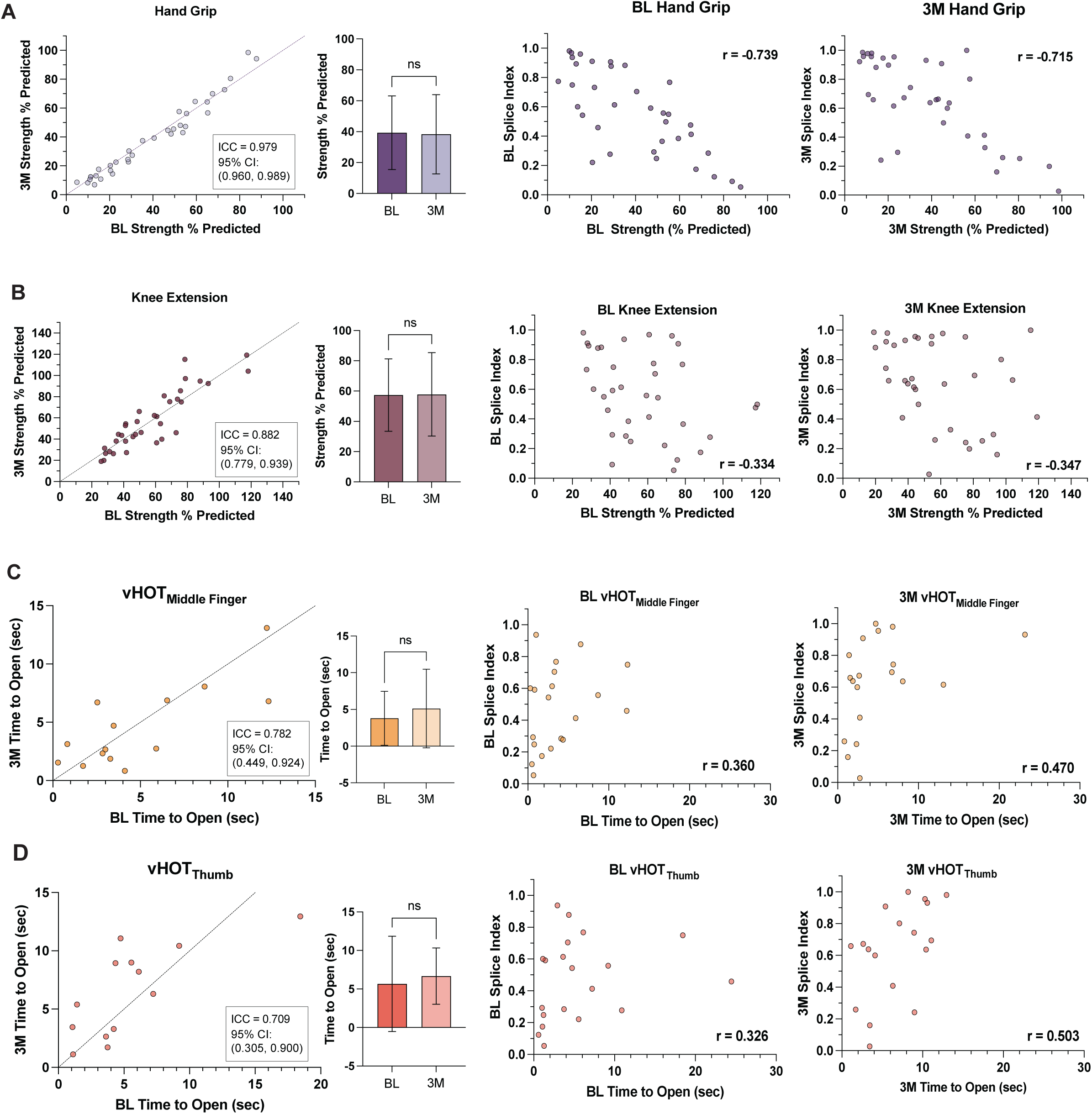
Baseline and 3-month SI score correlations to timepoint matched muscle strength and myotonia outcome measures in longitudinal DM1 sub-cohort. Correlation plot of baseline (BL) versus 3-month (3M) outcome measures in longitudinally sampled DM1 cohort. Line of agreement (x = y) and ICC with 95% CI are displayed. Bar plot of mean outcomes at BL and 3M is also shown for each outcome measure. Data presented as mean ± SD; paired t-test, ns = not significant. Correlation plots of BL SI and 3M SI versus timepoint matched measures are also displayed for each assessment. All correlations are reported as Pearson or Spearman r [95% CI]. a) Hand grip strength (HGS). Individual measures are reported as the percent of predicted strength as compared to unaffected individuals. BL SI v. BL HGS Pearson *r* = -0.739 [-0.860, -0.538], p < 0.0001, n = 35 & 3M SI v. 3M HGS Pearson *r* = -0.715 [-0.846, -0.504], p < 0.0001, n = 35 b) Knee extension (KE). Individual measures are reported as the percent of predicted strength as compared to unaffected individuals. BL SI v. BL KE Pearson *r* = -0.334 [-0.6002, --0.0005], p = 0.0501, n = 35 & 3M SI v. 3M KE Pearson *r* = -0.347 [-0.610, -0.016], p = 0.0411, n = 35. c) Video hand opening time (vHOT) of the middle finger. Individual measures are reported as time to open closed fist (seconds). BL SI v. BL vHOT_Middle Finger_ Spearman *r* = 0.360 [-0.113, 0.699], p = 0.1196, n = 20 & 3M SI v. 3M vHOT_Middle Finger_ Spearman *r* = 0.470 [0.006,0.768], p = 0.042, n = 19. d) vHOT of the thumb. BL SI v. BL vHOT_Thumb_ Spearman *r* = 0.326 [-0.150, 0.680], p = 0.1603, n = 20 & 3M SI v. 3M vHOT_Thumb_ Spearman *r* = 0.503 [0.032, 0.791], p = 0.0335, n = 35.

**Supplemental Figure 8:**
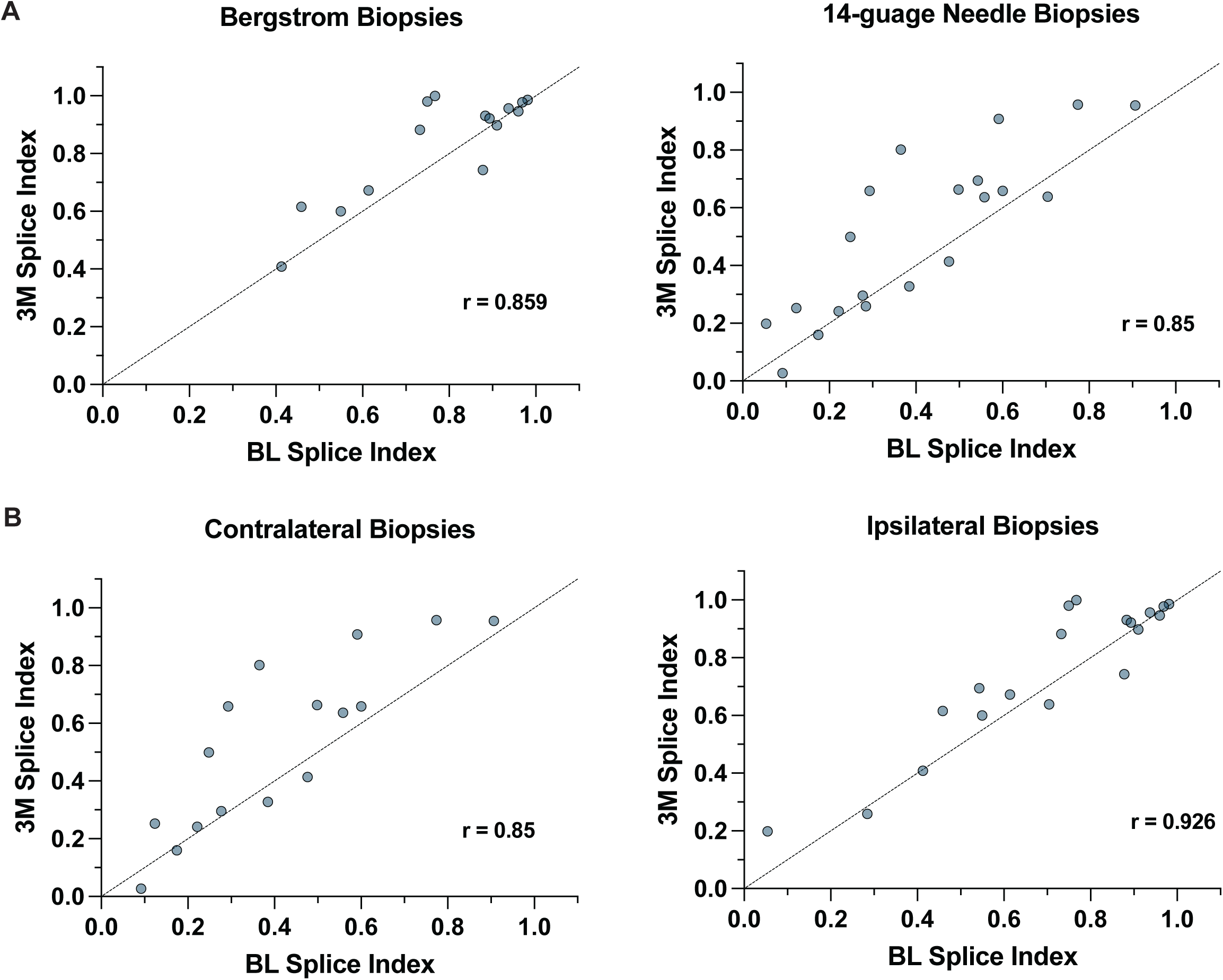
Correlation of baseline and 3-month SI scores in longitudinally sampled DM1 participants is similar independent of biopsy type or leg used for repeat biopsy. a) Correlation plot of BL versus 3M SI of longitudinally sampled DM1 subjects where muscle biopsies were collected via Bergstrom or 14-guage needle biopsy (Pearson *r* = 0.859 [0.618, 0.952], n = 15, p < 0.0001 & Pearson *r* = 0.85 [0.652, 0.939], n = 20, p < 0.0001, respectively). b) Correlation plot of BL versus 3M SI of longitudinally sampled DM1 subjects where 3M biopsy was collected from contralateral or ipsilateral tibialis anterior muscle (Pearson *r* = 0.85 [0.607, 0.946], n = 16, p < 0.0001 & Pearson *r* = 0.926 [0.813, 0.971], n = 19, p < 0.0001, respectively). All correlations are reported as Pearson r [95% CI]. Line of agreement (x = y) is displayed on all plots.\

**Supplemental Figure 9:**
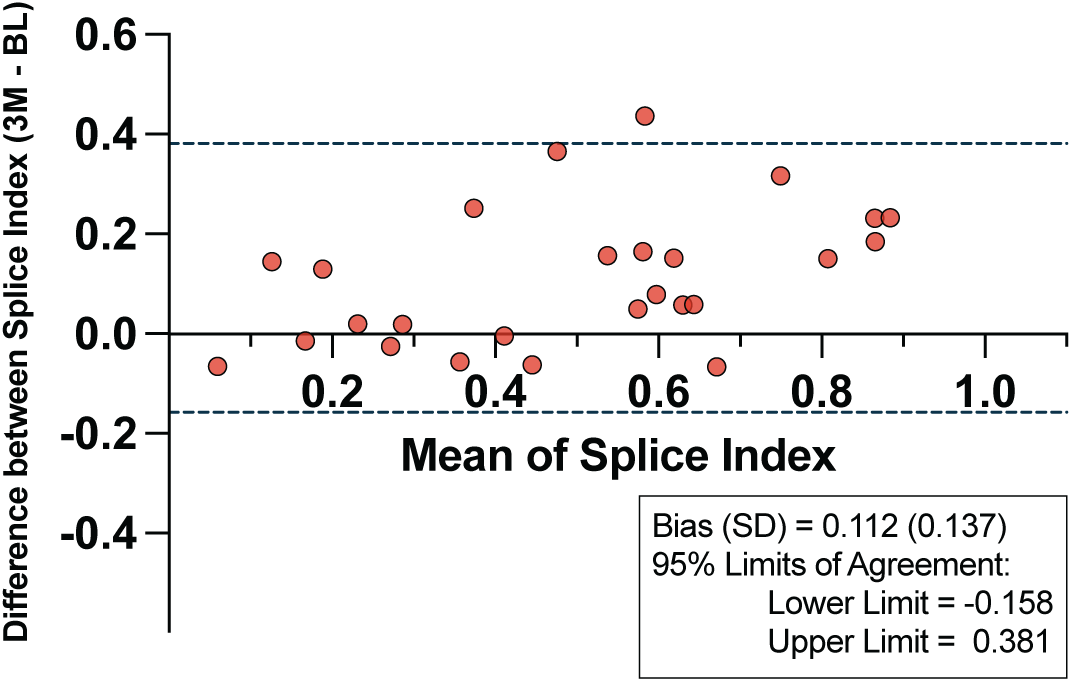
Participants with high SI scores anchor relative stability of SI between baseline and 3-months. Bland Altman plot illustrating agreement between Splicing Index scores at BL and 3M in DM1 longitudinal DM1 sub-cohort excluding individuals with high average SI scores (Mean SI > 0.8). Bias ± SD with 95% CI are displayed. ICC = 0.776, 95% CI [0.326, 0.914], p = 0.00175, n = 26.

**Supplemental Figure 10:**
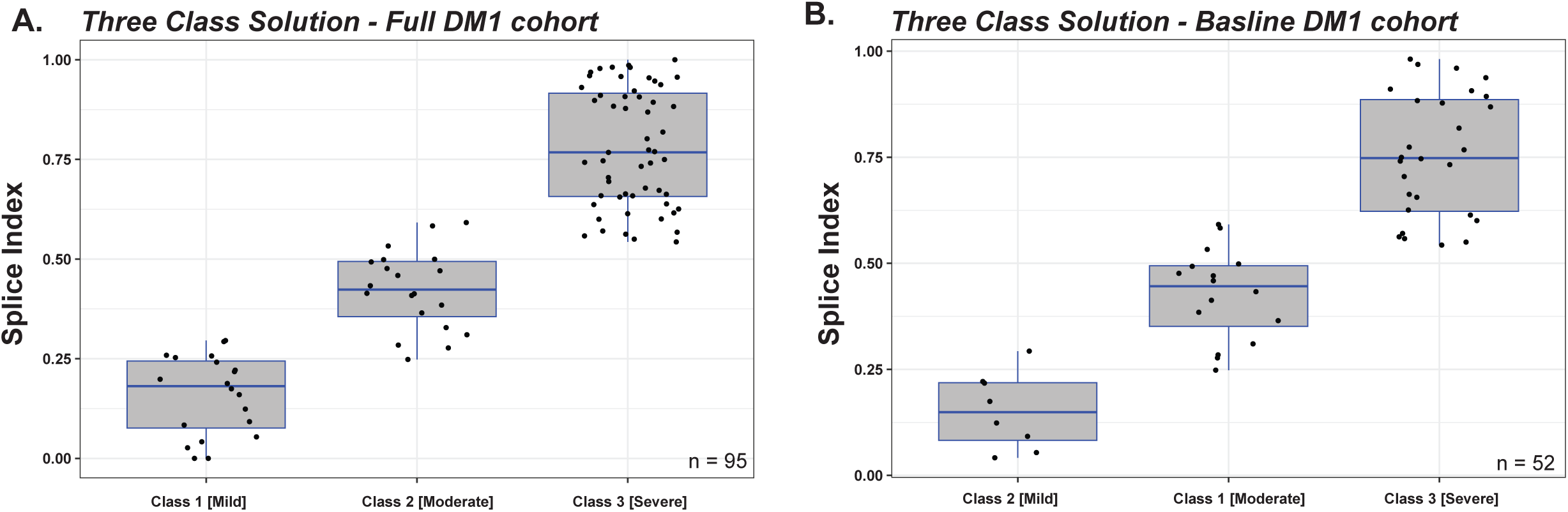
Distribution of Splicing Index values by latent class assignment. Box and whisker plot of splicing index (SI) values in latent class assignments. Line represents median value and whiskers extend to minimum and maximum value. The points show actual participant SI values. Random noise has been added to the horizontal position (i.e., x-axis) but not the vertical position (i.e., y-axis) to reduce overplotting. Statistical analyses of each LCA model and comparison of class demographics and associated outcome measures are reported in Sup Table 8. (a) LCA class assignment derived using entire study cohort with available targeted RNAseq data (n = 129). Only DM1 participants are plotted (n = 95). (b) LCA class assignment derived using DM1 participants with baseline functional assessments only (n = 52).

**Supplemental Figure 11:**
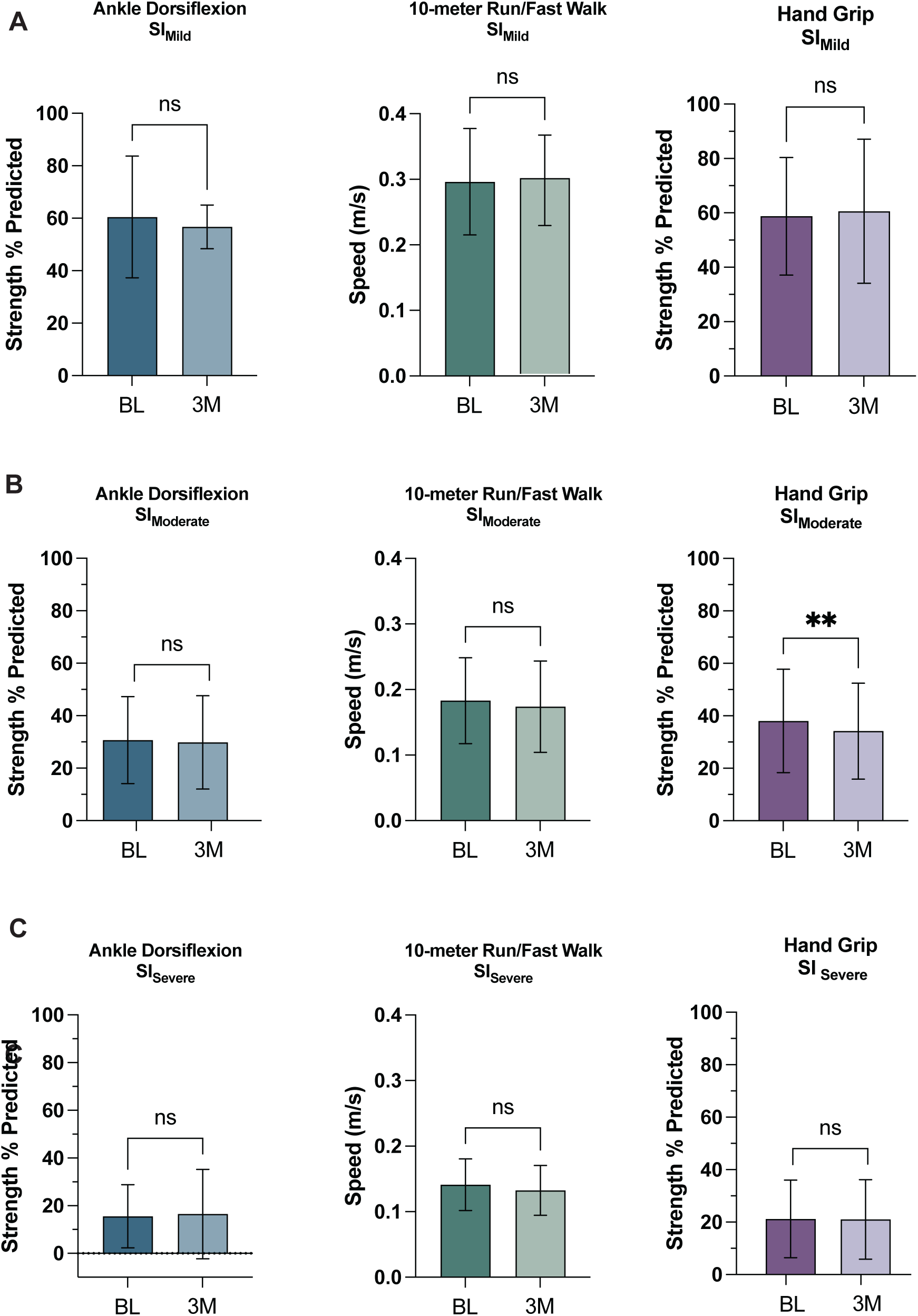
Mean performance on muscle strength and functional motor assessments are unchanged between BL and 3M timepoints in SI stratified sub-cohorts for nearly all clinical outcomes. a-c) Bar plot of ankle dorsiflexion (ADF) % predicted, 10-meter run/fast walk speed (m/s), and hand grip strength (HGS) % predicted at baseline and 3-months in baseline SI stratified sub-cohorts - SI_Mild_, SI_Moderate_, and SI_Severe_, respectively. Data represented as mean ± SD; paired t-test, ns = not significant, **p < 0.001

**Supplemental Figure 12:**
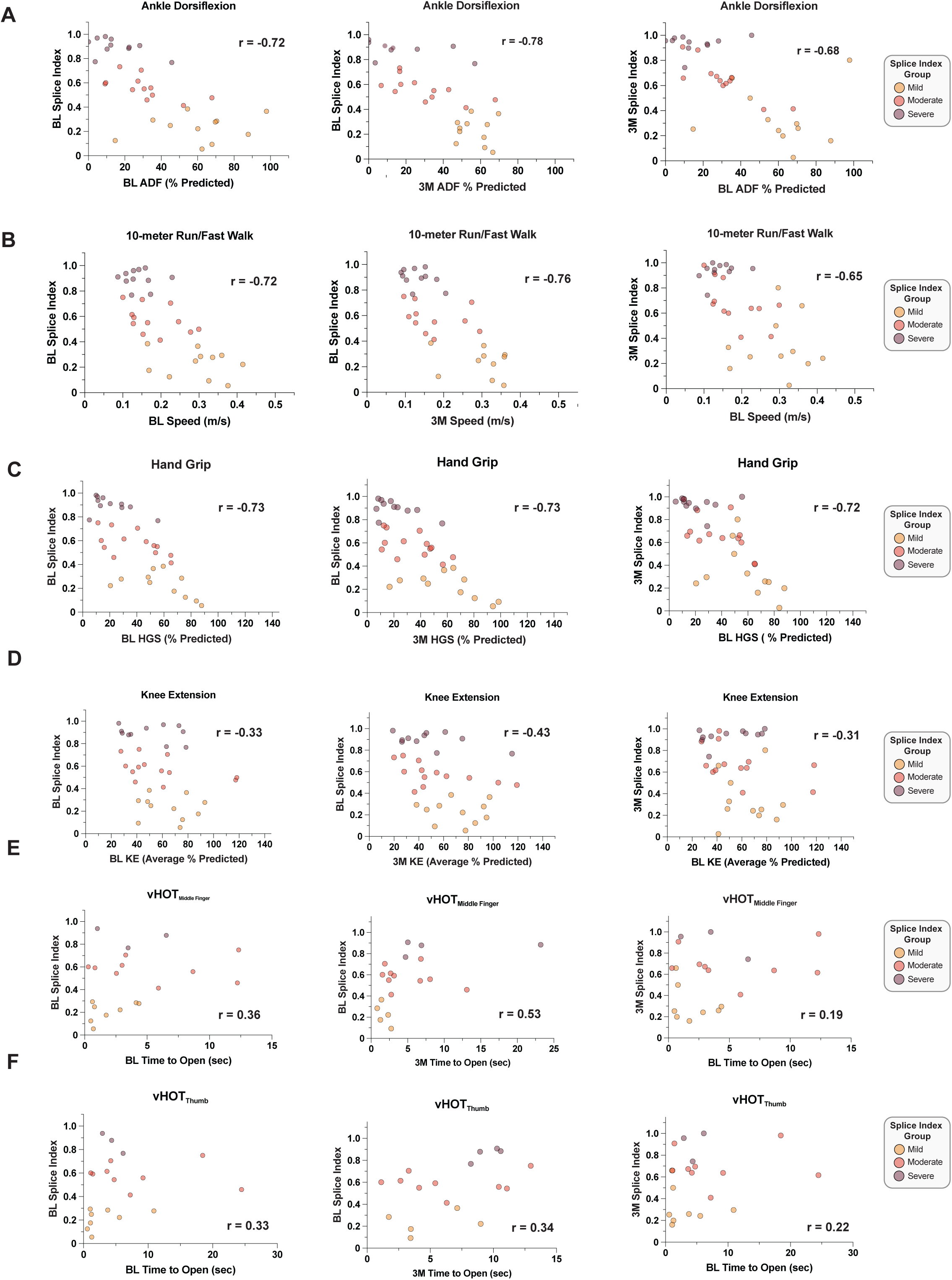
Timepoint mis-match correlation analysis of baseline and 3-month SI scores indicate potential predictive utility of the SI. Timepoint matched and mis-matched correlation plots of BL and 3M SI from the DM1 longitudinal cohort versus muscle strength, ambulation, and myotonia outcome measures at both timepoints. From left to right: BL SI v. BL measure, BL SI v. 3M measure, and 3M SI v. BL measures. Individual samples are colored by Mild, Moderate, and Severe sub-cohorts. All correlations are reported as Pearson or Spearman r [95% CI]. a) Ankle dorsiflexion (ADF). Individual measures are reported as the percent of predicted strength as compared to unaffected individuals. BL SI v. BL ADF showed a Pearson correlation coefficient of -0.7215 (95% CI = [-0.8518, -0.5070], p < 0.0001, n = 34). BL SI v. 3M ADF had a Pearson correlation coefficient of -0.7796 (95% CI = [-0.8846, -0.5996], p < 0.0001, n = 34). 3M SI v. BL ADF showed a Pearson correlation coefficient of -0.6835 (95% CI = [-0.8299, -0.4492], p < 0.0001, n = 34). b) 10-meter run/fast walk (10MRW). Individual measures are reported as speed (m/s). BL SI v. BL 10MRW showed a Pearson correlation coefficient of -0.7253 (95% CI = [-0.8540, -0.5129], p < 0.0001, n = 34). BL Splice Index v. 3M 10MRW had a Pearson correlation coefficient of -0.7589 (95% CI = [-0.8758, -0.5579], p < 0.0001, n = 32). 3M SI v. BL 10MRW had a Pearson correlation coefficient of -0.6463 (95% CI = [-0.8079, - 0.3944], p < 0.0001, n = 34). c) Hand grip strength (HGS). Individual measures are reported as the percent of predicted strength as compared to unaffected individuals. BL SI v. BL HGS showed a Pearson correlation coefficient of -0.739 (95% CI = [-0.860, -0.538], p < 0.0001, n = 35). BL SI v. 3M HGS had a Pearson correlation coefficient of -0.727 (95% CI = [-08535, -0.5196], p < 0.0001, n = 35). Additionally, 3M SI v. BL HGS showed a Pearson correlation coefficient of -0.7149 (95% CI = [-0.8465, -0.5010], p < 0.0001, n = 35). d) Knee extension strength (KE). Individual measures are reported as the percent of predicted strength as compared to unaffected individuals. BL SI v. BL KE showed a Pearson correlation coefficient of -0.3337 (95% CI = [-0.6002, -0.0005083], p = 0.0501, n = 35). BL SI v. 3M KE had a Pearson correlation coefficient of - 0.4277 (95% CI = [-0.6660, -0.1101], P = 0.0104, n = 35). 3M SI v. BL KE showed a Pearson correlation coefficient of -0.3066 (95% CI = [-0.5805, 0.02967], p = 0.0732, n = 35). e) Video hand opening time (vHOT) of the middle finger. Individual measures are reported as time to open closed fist (seconds). BL SI v. BL vHOT_Middle Finger_ showed a Spearman correlation coefficient of 0.3594 (95% CI = [-0.1127, 0.6991], p = 0.1196, n = 20). BL SI v. 3M vHOT_Middle Finger_ had a Spearman correlation coefficient of 0.222 (95% CI = [-0.257, 0.691], p = 0.346, n = 20). Additionally, 3M SI v. BL vHOT_Middle Finger_ showed a Spearman correlation coefficient of 0.187 (95% CI = [0.292, 0.590], p = 0.4312, n = 20). f) Video hand opening time (vHOT) of the thumb. BL SI v. BL vHOT_Thumb_ showed a Spearman correlation coefficient of 0.3263 (95% CI = [-0.1496, 0.6795], p = 0.1603, n = 20). BL SI v. 3M vHOT_Thumb_ had a Spearman correlation coefficient of 0.316 (95% CI = [-0.164, 0.705], p = 0.165, n = 18). Additionally, 3M SI v. BL vHOT_Thumb_ showed a Spearman correlation coefficient of 0.592 (95% CI = [0.850, 0.798], p = 0.0196, n = 19).

**Supplemental Figure 13:**
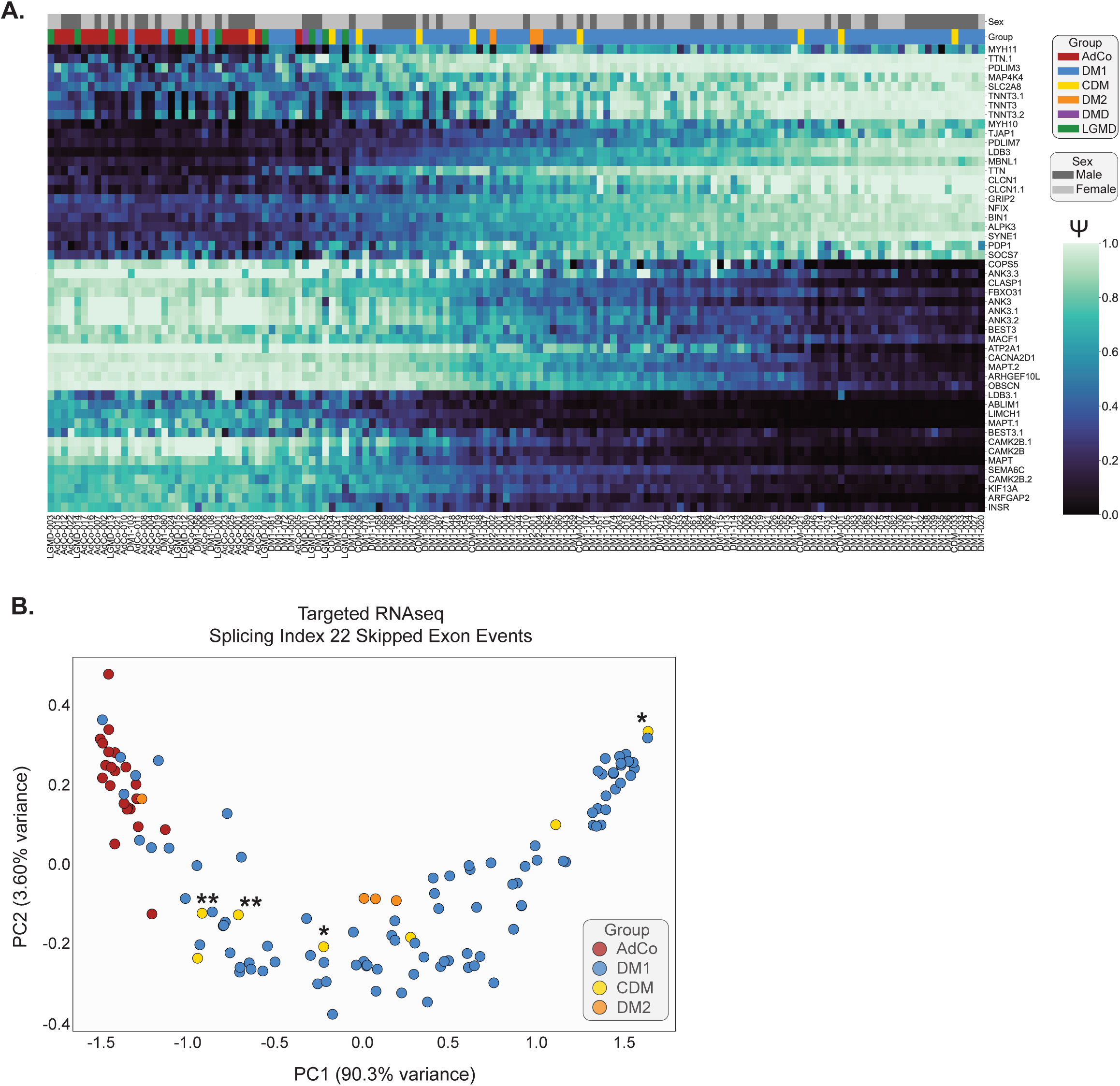
Splicing index captures splicing dysregulation in ambulatory DM2 subjects and longitudinally sampled CDM individuals. a) Heatmap displaying estimated percent spliced in (Ψ) of top 50 significantly dysregulated skipped exon (SE) events between all DM participants (DM1, DM2, and CDM) versus unaffected adult controls (AdCo) & disease control reference groups (DMD & LGMD) subjected to total RNAseq (|ΔΨ| ≥ 0.1, FDR ≤ 0.05) (Sup Table 10). Both rows (SE events) and columns (individual samples) were subjected to hierarchical clustering. Sample group and sex are annotated above the heatmap and individual Subject IDs are reported below. b) Principal component analysis of Ψ values for 22 splicing events encompassing the Splicing Index derived from targeted RNAseq inclusive of DM2 and CDM samples (Sup Table 5). Longitudinally sampled CDM participants are annotated (*CDM-001 and CDM-018 sampled at 2 weeks and 8 years of age; **CDM-032 and CDM-036 sampled at 12 and 16 years of age)

## Table Legends

<u>Supplemental Table 1:</u> *Demographic table of DM1 participants and other subjects*.

DM1 Participant ID, Subject ID, sex, age at biopsy, group, biopsy method, muscle biopsied, and contralateral/ipsilateral biopsy collection are reported. For each participant SI scores are reported at each timepoint (baseline or 3-months). Global spliceopathy measures derived from total RNA-seq are also listed (Mean ΔΨ). Where available, clinical measures of muscle strength, motor function, and vHOT are provided. SRA accession numbers of associated total and targeted RNAseq are also provided.

<u>Supplemental Table 2:</u> *Significantly dysregulated splicing events in DM1 skeletal muscle as determined by total RNAseq*.

Ψ values of 946 skipped exon events defined as significantly dysregulated between DM1 subjects versus unaffected adult controls (AdCo) and disease controls (LGMD, DMD) (|ΔΨ| ≥ 0.1, FDR ≤ 0.05). Events are rank ordered by mean inclusion level difference and Ψ values for each individual sample are listed.

<u>Supplemental Table 3:</u> *Ψ values and isoform counts of 22 skipped exon events encompassing Splicing Index panel derived from total RNAseq*.

Ψ values for all DM1, AdCo, LGMD, and DMD subjects subjected to total RNAseq are listed. Skipped exon event coordinates, median Ψ & 95% CI for each sample group, and event inclusion & exclusion isoform counts per subject are reported.

<u>Supplemental Table 4:</u> *Curve fitting parameters of Splicing Index events*.

Parameters derived from curve fitting of total RNAseq Ψ versus normalized mean ΔΨ using a four-parameter dose response curve (Fig 2A) are reported, including Ψ_min_, Ψ_max_, span, slope, EC_50_, and R^2^ of fit are listed. All values are reported as mean ± SEM. Pearson correlation with normalized mean ΔΨ for each event is also reported with the 95% confidence interval and associated p-value.

<u>Supplemental Table 5:</u> *Ψ values and isoform counts derived from targeted RNAseq of 22 skipped exon events encompassing Splicing Index*.

Ψ for all DM1, AdCo, CDM, and DM2 subjects subjected to targeted RNAseq of 22 event SI panel. Median Ψ & 95% CI for each sample group and event inclusion & exclusion isoform counts per subject are listed. Normalized Ψ values scaled using Ψ_Median Control_ and Ψ_DM95_ reference values (Sup Table 6) are also reported.

<u>Supplemental Table 6:</u> *Normative Ψ reference values derived for scaling of individual sample Ψ for 22 splice events within the composite Splicing Index*.

Ψ_Median Control_ and Ψ_DM95_ reference values derived from 22 unaffected adult controls (AdCo) and all 95 DM1 samples subjected to targeted RNAseq from the assembled cohort.

<u>Supplemental Table 7:</u> *Individual splice event associations with clinical outcome measures in complete cross-sectional DM1 cohort*.

Individual correlations of normalized targeted RNAseq Ψ (Sup Table 5) for all 22 splice events encompassed within the composite Splicing Index with outcomes assessments, including ADF, HGS, KE, 10-meter run/fast walk, vHOT_Thumb_, and vHOT_Middle Finger_. Number of matched samples (n), 95% CI, R^2^, and p-value are reported for each splice event. Events are listed and force-ranked by Pearson *r*.

<u>Supplemental Table 8:</u> *Latent class assignment model information and individual class correlations with functional outcome measures*.

Statistical measures of LCA models goodness of fit using two and three class solutions in entire study cohort with targeted RNAseq data (full cohort, n = 129) and DM1 subjects with baseline outcome measures (BL only, n = 52). Comparison of demographic variables (age, sex) and mean functional outcomes measures between LCA defined classes is also reported.

<u>Supplemental Table 9:</u> *Complete intercept table of multiple linear regression Models 1-3*. Companion table to regression models reported in Figure 7 & Table 1. Companion table to regression models reported in Figure 7 & Table 1. Multiple linear regression analysis evaluating the effects of baseline Splice Index (BL SI) and baseline ADF on the dependent variable of ADF performance (3M ADF) or progression from baseline [ Δ ADF(3M-BL) ]. The analysis includes an examination of variance components, parameter estimates, and goodness-of-fit metrics. Multiple statistical elements are reported (sum of squares (SS), degrees of freedom (DF), mean squares (MS), F-statistic (F), standard errors (SE), 95% confidence intervals (CI), t-values, and p values.

<u>Supplemental Table 10:</u> *Top 50 most significantly dysregulated RNA splicing events as in all DM1, DM2, and CDM individuals as determined by total RNAseq.* Ψ values of the top 50 most significantly dysregulated skipped exon events between DM1, CDM, and DM2 subjects versus unaffected adult controls (AdCo) & disease controls (LGMD, DMD) (|ΔΨ| ≥ 0.1, FDR ≤ 0.05). Events are rank ordered by mean inclusion level difference.

<u>Supplemental Table 11:</u> *Splicing Index reference materials.* Table containing hg38 genome coordinates for 22 skipped exon events encompassed within the Splicing index, cassette exon sequence, multiplex PCR primers and cycling conditions, amplicon information and sequence, and custom reference sequence set for derivation of exon inclusion and exclusion counts.

